# Genetically engineered retina for improved retinal reconstruction after transplantation

**DOI:** 10.1101/2020.12.23.424068

**Authors:** Take Matsuyama, Hung-Ya Tu, Jianan Sun, Tomoyo Hashiguchi, Ryutaro Akiba, Junki Sho, Momo Fujii, Akishi Onishi, Masayo Takahashi, Michiko Mandai

**Author notes:** These authors contributed equally (Take Matsuyama, Hung-Ya Tu).

## Abstract

ES/iPS-retinal sheet transplantation, which supplies photoreceptors as well as other retinal cells, has been shown able to restore visual function in mice with end-stage retinal degeneration. Here, by introducing a novel type of genetically engineered ES/iPS-retinal sheet with reduced numbers of secondary retinal neurons but intact photoreceptor cell layer structure, we reinforced the evidence that ES/iPS-retinal sheet transplantation can establish synaptic connections with the host, restore light responsiveness and reduce aberrant RGC spiking. Furthermore, we show that genetically engineered grafts can substantially improve the outcome of the treatment by improving neural integration. We speculate that this leads to reduced spontaneous activity in the host which in turn contributes to a better visual recovery.

## Introduction

Several cell-based therapies are currently under development to treat photoreceptor cell loss^1–3^. While transplantations may be conducted on hosts with varying degrees of retinal degeneration using donor or pluripotent stem cell derived materials, the overall strategy can be categorized into either single cell photoreceptor suspensions or retinal sheet transplantation. Cell suspension strategies consist of transplanting purified photoreceptor precursor cells^4–6^, whereas retinal sheet transplantations engraft retinal organoids containing both photoreceptor cells and inner retinal neurons^7–9^. Both methods have shown cellular integration, maturation of the photoreceptor cells and some recovery of light responsiveness^10–15^.

The inner neurons in the retinal sheet transplantation can serve as a scaffold and nurturing microenvironment for photoreceptor cells to differentiate and mature, producing an organized layered structure resembling the retina and well-developed photoreceptor morphology^8,9,12^, whereas single cell photoreceptor transplantation seldom form organized layered structures or outer segments. On the other hand, single cell photoreceptor suspension preparations may have a better chance to contact host cells with directly exposed photoreceptor cells to host bipolar cells, whereas inner neurons in retinal sheet transplantation may become a hindrance for graft-host neural integration, as graft photoreceptors form synapses with concomitantly developing bipolar cells, resulting in numerous intra-graft synapses.

In this study, we attempted to bring the best of the two approaches together by using genetically engineered cell lines to prepare retinal sheets with fewer bipolar cells. To this end we prepared two cell lines, *Bhlhb4^-/-^* and *Islet1^-/-^*. The basic helix-loop-helix family member e23 (*Bhlhe23*, also known as *Bhlhb4*) is a transcription factor^16^ expressed in rod bipolar cells, and it is required for maintenance and neural maturation of rod bipolar cells^17^. *Bhlhb4* functions downstream of factors that determine bipolar cell specification, and thus *Bhlhb4* retinas can engage rod bipolar cell fate, however, nascent cells that fail to mature eventually succumb to apoptosis^17^. *Islet1* (or *Isl1*), a LIM-homeodomain transcription factor mediates neuronal differentiation of rod and cone bipolar, cholinergic amacrine, and ganglion cells in the retina. Deletion of *Islet1* results in the loss of the majority of bipolar, cholinergic amacrine, and ganglion cells^18^. Due to the relatively late expression of *Bhhb4* and *Islet1*, we speculated that deletion of these genes would not largely affect retinal development before bipolar differentiation, but grafts would have fewer bipolar cells around the time photoreceptors start to form synapses. We prepared retinal sheets from wildtype (wt), *Bhlhb4^-/-^*, and *Islet1^-/-^* pluripotent cell lines (both iPS and ES) and transplanted them to end-stage *rd1* mouse retinas, in which rod photoreceptors are virtually abolished by postnatal day 30 and surviving cone photoreceptors degenerate secondarily. These genetically engineered cell lines can be differentiated to retinal organoids with similar potency as wt cell lines and integrate with host retinas after transplantation. Our analyses of post-transplantation retinas show that *Bhlhb4^-/-^* and *Islet1^-/-^* result in a reduced bipolar cell population and a concomitant improved neural integration. We further conducted detailed electrophysiological analyses that indicate the improved neural integration may be manifested as a decrease in spontaneous activity, which in turn is observed as an enhanced performance in behavior tests. Overall our results show that *Bhlhb4^-/-^* and *Islet1^-/-^* can be used to improve neural integration of retinal sheet transplantation.

## Results

### Bhlhb4^-/-^ and Islet1^-/-^ cells differentiate to retinal organoids similarly to wt

Both *Bhlhb4^-/-^* and *Islet1^-/-^* cell lines were able to differentiate into retinal structures similarly to wt cell lines. All the *Bhlhb4^-/-^* (6 clones) and *Islet1^-/-^* (4 clones) cells (iPS and ES cells) self-reorganized to homogenous aggregates and formed hollowed structures resembling the optic vesicle by differentiation day (DD) 7. These retinal organoids grew bigger by sprouting more vesicular structures through DD23 (Figure 1a). Transgenic cell lines expressing GFP under the *Nrl* promoter, which is specifically activated in rod photoreceptors shortly after terminal cell division^19^, started to show signs of detectable fluorescence around DD19. The GFP signal consistently grew stronger until DD33 and then the fluorescence became weaker. The *Nrl*-GFP fluorescence signal between DD19 and DD33 of representative clones was modeled with an exponential growth curve revealing that *Bhlhb4^-/-^* and *Islet1^-/-^* cell lines have broadly similar time courses and maximum intensities to wt lines (Figures 1b, 1c and Supplemental Figure 1), indicating that these gene knockouts do not affect early retinal differentiation.

**Figure 1.**
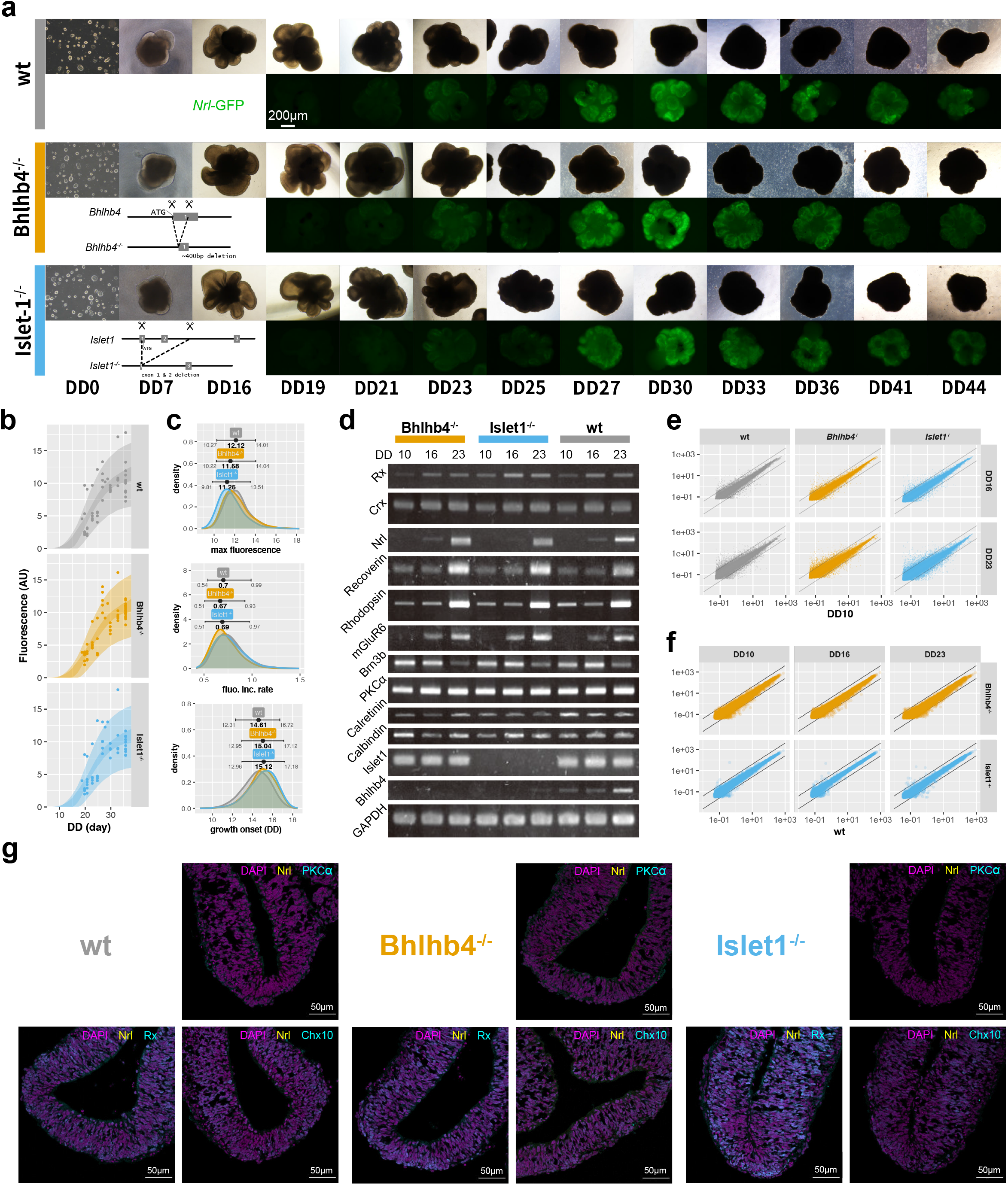
Differentiation of retinal organoids. (a) Phase contrast and fluorescence images of differentiating retinal organoids at different differentiation days (DD). (b) Modeling of *Nrl*-GFP signal with a growth curve. Points represent the intensity value for individual retinal organoids, the light shaded area shows the range of expected values, and the dark shaded area shows the expected mean signal intensity with 89% compatibility intervals. Full parameter estimates for the model are provided in Supplemental Figure 1. (c) Estimated parameter values for maximum fluorescence value (top), maximum rate of fluorescence increase (middle), and onset of fluorescence increase (bottom). Estimates largely overlap showing no differences among organoids. n=170 organoids (54 wt, 61 *Bhlhb4^-/-^*, and 55 *Islet1^-/-^*) were used for this analysis. (d) RT-PCR of key genes from organoids differentiated from a clone of Tg(*Nrl*-GFP) iPS cell lines. Similar results were obtained for ES derived organoids (Supplemental Figure 1). (e-f) Scatter plots of microarray expression pattern of retinal organoids shows no large differences among retinal organoids. (e) shows the gene expression of DD16 and DD23 compared to DD10 and (f) shows the comparison between *Bhlhb4^-/-^* and Iselt1^*-/-*^ to wt. Diagonal lines correspond to 4-fold expression differences. GO analysis of differentially expressed genes is provided in Supplemental Figure 2. (g) Immunohistochemistry characterization of PKCα, Chx10, and Rx expression in wt, *Bhlhb4^-/-^* and *Islet1^-/-^* grafts on DD15.

*Bhlhb4* expression begins around postnatal day 5 (P5) in mouse retinas, which is roughly equivalent to DD25 of the retinal organoids and is restricted to developing rod bipolar cells^17^. Consistently, we observed *Bhlhb4* mRNA expression gradually increase on wt and *Islet1^-/-^* retinal organoids from DD10 to DD23, whereas the band was absent on *Bhlhb4^-/-^* organoids (Figure 1d and Supplemental Figure 1). Similarly, *Islet1* expression in bipolar cells begins around P5, however *Islet1* is also expressed at earlier stages in the RGC^18^. Consequently, *Islet1* is detected on DD10, DD16, and DD23 in wt and *Bhlhb4^-/-^* retinal organoids but is absent in *Islet1^-/-^* (Figure 1d and Supplemental Figure 1). We saw no differences between ES and iPS derived organoids in the expression pattern of key genes, and the effect of *Bhlhb4* or *Islet1* deletion was also identical (Figure 1d and Supplemental Figure 1). In addition to RT-PCR, we conducted the microarray analyses of DD10, DD16, and DD23 organoids derived from *Nrl*-GFP;Ribeye-reporter iPSC lines. Supplemental Figure 1 shows some key genes specific to retinal progenitors, bipolar, and photoreceptor cells. The expression pattern of these genes is largely unaltered. Although *Bhlhb4^-/-^* retinal organoids seem to have lower expression of some photoreceptor markers (*Cnga1, Cnga2, Cnga3, Gucy2d, Rho*), and *Islet1^-/-^* seem to have higher expression (*Cabp4, Cnga1, Cngb1, Gnat2, Gngt1, Rho, Rcvrn, Sag*), photoreceptor markers are generally low at this early developmental stage. Expression patterns for RPE, glial cells, retinal ganglion cells, amacrine cells, horizontal cells, as well as other controls (pluripotency, neuron, apoptosis, and expression reference marker genes) were also largely identical amongst the three lines. Overall gene expression across cell lines was very similar, with few genes up-regulated or down-regulated compared to the wt line (Figures 1e and 1f). Gene Ontology (GO) enrichment analysis shows similar genes, mostly pertaining to localization, metabolic, and developmental processes, are up-regulated or down-regulated across development (Supplemental Figure 2).

Figure 1g shows immunostaining for *PKCα, Chx10*, and *Rx*, as well as the *Nrl*-GFP reporter fluorescence of DD15 retinal organoids *in vitro*. At DD15, a widespread expression of *Rx* is observed reflecting the early stage of differentiation. Virtually no *PKCα* or *Nrl* expression is evident at this stage, although *Chx10*, which is expressed in some progenitor cells, is weakly expressed in some cells. The same trend is observed on wt, *Bhlhb4^-/-^*, and *Islet1^-/-^* graft lines. Overall our data shows that *Bhlhb4^-/-^* and *Islet1^-/-^* cells can differentiate to retinal organoids without any apparent differences to wt cells.

### Bipolar cells are markedly reduced in Bhlhb4^-/-^ and Islet1^-/-^ grafts after transplantation

We transplanted early developmental stage (DD10-DD15) retinal organoids to the subretinal space of *rd1* mouse and observed maturation and integration with the host. Grafts were examined after 4-5 weeks after transplantation, that is, 6-7 weeks old grafts. Transplanted grafts consistently developed photoreceptor layers with typical small and dense nucleus and strong Rhodopsin expression at segment-like structures inside the rosettes, surrounded by inner retinal cells (Figure 2c and Supplemental Figure 3a). We first assessed if PKCα^+^ rod bipolar cells were actually reduced as had been primarily aimed in this study. In grafts derived from all the *Bhlhb4^-/-^* and *Islet1^-/-^* clones, the number of PKCα^+^ cells were consistently and markedly reduced (Figures 2a and 2b). Our data shows that PKCα^+^ cells account for about 0.14[89%CI 0.05-0.34] of the inner cells in wt and about 0.04[89%CI 0.01-0.12] and 0.02[89%CI 0.01-0.07] in *Bhlhb4^-/-^* and *Islet1^-/-^* grafts respectively (confidence in the difference in the effect of graft lines i.e., the fraction of area over zero of the difference in the distributions, were 97% for *Bhlhb4^-/-^*-*wt*, 99% for *Islet1^-/-^*-*wt*, and 84% for *Bhlhb4^-/-^-Islet1^-/-^*). On the other hand, cone bipolar cells (SCGN^+^) seemed to decrease slightly in *Islet1^-/-^* (0.19[89%CI 0.11-0.27]) compared to the wt (0.28[89%CI 0.19-0.39]), whereas the decrease in *Bhlhb4^-/-^*(0.25[89%CI 0.16-0.33]) is not as evident (confidence in the difference in the effect of graft lines for SCGN^+^ cells were 72% for *Bhlhb4^-/-^-wt*, 93% for *Islet1^-/-^-wt*, and 86% for *Islet1^-/-^-Bhlhb4^-/-^*). These results are consistent with the notion that *Bhlhb4* affects rod bipolar cells specifically whereas *Islet1* affects both rod and cone bipolar cells. Pan-bipolar marker Chx10^+^ cells were also reduced in *Bhlhb4^-/-^* and slightly less in *Islet1^-/-^* (confidence in the difference in the effect of graft lines for Chx10^+^ cells were 99% for *Bhlhb4^-/-^-wt*, 84% for *Islet1^-/-^-wt*, and 99% for *Bhlhb4^-/-^-Islet1^-/-^*). It is noteworthy to mention that Chx10^+^ cells in *Islet1^-/-^* were characterized by a weaker signal, indicating these cells were different from typical bipolar cells. The number of Calbindin^+^ and Calretinin^+^ cells were similar among wt, *Bhlhb4^-/-^, Islet1^-/-^* (Supplemental Figure 3). Additionally, we investigated if photoreceptor population was changed in *Bhlhb4^-/-^* and *Islet1^-/-^* grafts due to the reduction of the number of bipolar cells. The size of rosettes varies considerably, but the number of rod photoreceptor cells seems to be proportional to the number of graft cells (Figures 2c and 2d). We found no significant differences in the photoreceptor content, with all the lines producing grafts of about 0.6 rod photoreceptor content (Figure 2d and Supplemental Figure 3). Finally, while most of the grafted cells formed spherical rosette structures, substantial organization/reconstruction of highly polarized ONL structure was observed in some cases (Figure 3a, *Bhlhb4^-/-^*). This was sporadically observed and seemed to be independent of the graft genotype (wt, *Bhlhb4^-/-^, Iselet1^-/-^*). Although we do not yet know the specific conditions leading to these cases, they showcase the regenerative potential after transplantation in the end-stage degenerate retina.

**Figure 2.**
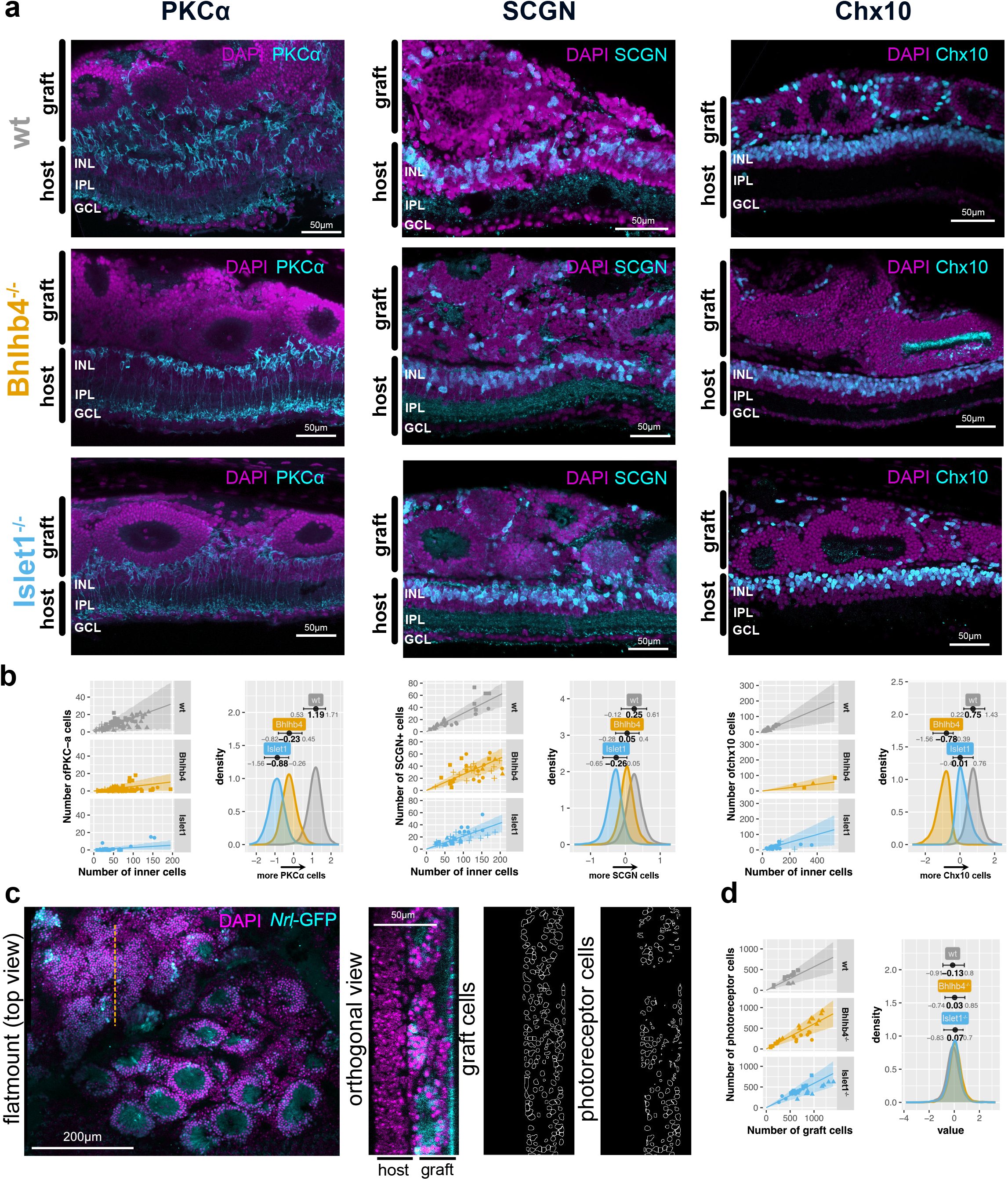
Immunohistochemistry characterization of wt and *Bhlhb4^-/-^* and *Islet1^-/-^* grafts after subretinal transplantation. (a) PKCα^+^ (rod bipolar), SCGN^+^ (cone bipolar), and Chx10^+^ (pan-bipolar) cells in wt, *Bhlb4^-/-^*, and *Islet1^-/-^* grafts. (b) Quantification of cell markers. Left plots show the number of PKCα^+^, SCGN^+^, and Chx10^+^ cells (vertical axis) against the number of inner cells in the graft per section (horizontal axis). Dots indicate the count value for a given section, with different shapes representing samples from different animals. The line represents the most likely value and the shaded area represents the 89% compatibility interval assuming that the number of positive cells for a positive marker follows the binomial distribution given the number of total (inner) cells. Right plots show the estimated effect of graft genotype on the probability of the marker being positive, with mode and 89% compatibility intervals indicated on top. Note that the horizontal scale (value) is logit, i.e. log odds, with higher values indicating higher probability of observing a cell positive for a particular marker. Distribution for the rest of the parameters, as well as results for a similar analysis on Calbindin^+^ and Calretinin^+^ cells are provided in Supplemental Figure 2. (c) Flat mounted retinas were imaged and orthogonal sections intersecting rosettes were constructed to analyze the number of photoreceptor cells. The images to the right correspond to orthogonal views at the dotted orange line. (d) The number of photoreceptor cells (vertical axis) was modeled as a function of number of graft cells (horizontal axis), similar to (b). The graph on the right shows there is no difference in photoreceptor ratio by graft genotype as the posterior distributions largely overlap. Distribution for the rest of the parameters is provided in Supplemental Figure 2 as well as predicted values from the model. 11 mice were used for PKCα^+^ analysis (3 wt, 3 *Bhlhb4^-/-^*, and 4 *Islet1^-/-^*), 11 for SCGN^+^ analysis (3 wt, 4 *Bhlhb4^-/-^*, and 4 *Islet1^-/-^*), 9 for Chx10^+^ cells (2 wt, 3 *Bhlhb4^-/-^*, and 4 *Islet1^-/-^*), and 8 for the photoreceptor cell number analysis (2 wt, 3 *Bhlhb4^-/-^*, and 3 *Islet1^-/-^*). GCL: ganglion cell layer, INL: inner nuclear layer, ONL: outer nuclear layer.

**Figure 3.**
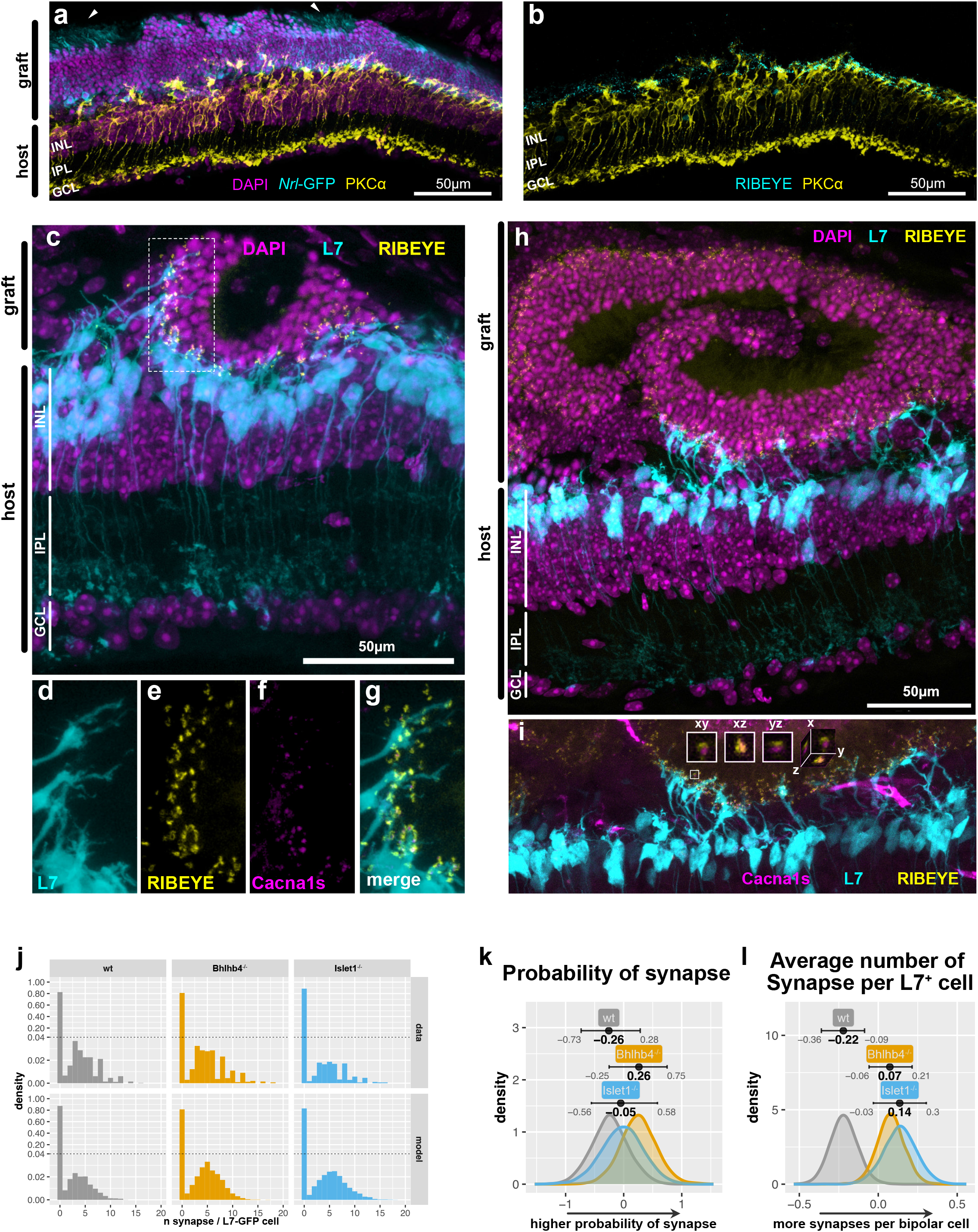
Host-graft synapse formation. (a,b) Example of graft contacting host forming an outer nuclear layer (ONL) reminiscent of wildtype retina. PKCα positive host bipolar cells extend dendrites with the presynaptic marker (RIBEYE) at their tips and graft ONL shows oriented IS/OS like structures (arrow heads). Representative images of synapse formation from *Bhlhb4^-/-^* grafts (c-g). (c) *L7*-GFP host rod bipolar cells (cyan) are extending dendrites toward graft photoreceptor cells. (d-g, and i) Example of synapse quantification. (d) Shows the *L7*-GFP host bipolar cell dendrites. RIBEYE (e) and Cacna1s (f) pairs at the tip of *L7*-GFP cells were treated as host-graft synapses (g). (h,i) Heterogeneity in synaptic integration. The right part of the image shows extensive synaptic contact, with *L7*-GFP bipolar aggressively extending towards graft photoreceptor cells while host bipolar cells on the left are completely retracted and show no synapse formation. (j) Histogram of number synapse per *L7*-GFP cell (host bipolar). Note that the scaling of the vertical axis has been altered to facilitate visualization (dotted line). Top panels represent the raw data while bottom rows represent the modeled data. Most of the host bipolar cells do not form synapses with the graft, resulting in a large zero-count. The effect of the graft on the probability of forming synapse (k) and the average number of synapses per L7-GFP cell (l), respectively, was estimated using a zero-inflated Poisson model and shown with 89% compatibility intervals and modes indicated. Note that the horizontal scale for (k) is in log odds and (l) in log. Larger values in (k) indicate larger probability of synapse and higher values in (l) indicate more synapses per bipolar cell. Complete description of the model parameters can be found in Supplemental Table 1 along with predicted values from the model. GCL: ganglion cell layer, INL: inner nuclear layer, ONL: outer nuclear layer. 10354 observation (L7-GFP bipolar cells) from 27 mice (9 wt, 9 *Bhlhb4^-/-^*, 9 *Islet1^-/-^*) were used for this analysis.

### Bhlhb4^-/-^ and Islet1^-/-^ grafts have more synapses per host bipolar cell

We have often seen images indicative of robust neural integration, such as in Figures 3a-b, with host bipolar dendrites extending towards graft cells and surrounded by photoreceptor presynaptic marker RIBEYE^20^. It is, however, difficult to confirm bona fide synapses formed between host bipolar cells and graft photoreceptor cells (hereinafter referred to as “*de novo*” synapse), as graft bipolar cells may also form synapses. In order to identify *de novo* synapses we used *rd1;L7*-GFP mice as host and grafts derived from the Ribeye-reporter cell line. Additionally, we immunostained the postsynaptic marker Cacna1s and counted synapses on the dendritic tips of host *L7*-GFP^+^ rod bipolar cells, ensuring that we are counting genuine synapses, and that these are between host and graft cells. Figures 3c-g show an example of host-graft synapse formation in *Bhlhb4^-/-^* line. *L7*-GFP^+^ bipolar cells forming *de novo* synapses exhibited the typical candelabra-like dendritic arbor of normal rod bipolar cells (Figure 3c, Supplemental Movie 1) in contrast to the dendrite-less morphology of the degenerate retina. Notably, host rod bipolar cells sometimes extended their dendrites far into the graft area (Figures 3h-i, right area). The majority of host bipolar cells do not make contact at all, but when they do, multiple synapses are formed between a host bipolar cell and graft photoreceptor cells, resulting in synapse distribution with a sharp zero count. Figure 3j shows the distribution of the number of synapses per host bipolar cell (*L7*-GFP^+^). This data is best described as a mixture of two distributions with two parameters, known as “zero inflated Poisson”. One distribution, the large zero counts, describes the probability of forming a synapse at all (*θ*), and the other one, the broad peak, describes the average number of synapses (*λ*) once synaptic contact is established (Supplemental Figure 4a). The probability of host bipolar cells to form synapses varied greatly (0.09-0.22), with no clear differences amongst the lines (Figure 3k and Supplemental Figure 4b; confidence in the difference in the effect of graft line was 83% for *wt-Bhlhb4^-/-^*, 65% for *wt-Islet1^-/-^*, and 68% for *Islet1^-/-^-Bhlhb4^-/-^*). There was, however, a notable increase in the average number of synapses per host bipolar cell in the KO lines (Figure 3l; confidence in the difference in effect of graft line was 98% for *Bhlhb4^-/-^-wt*, 98% for *Islet1^-/-^-wt*, and 64% for *Bhlhb4^-/-^-Islet1^-/-^*). Interestingly, we also found that sex of the host has a significant effect on the outcome resulting in more synapses per bipolar in females (Supplemental Figure 4c; confidence in the difference in the effect of sex was 91% for *male-female*). In addition to these main effects (overall effects), we estimated the interactions, conditional effects that are present under different combinations, of genotype and sex (Supplemental Figure 4, genotype x sex panels). Together the effects of line and sex and their interaction indicate that wt grafts produce 3-6 synapses (male:3.25[89%CI 2.54-4.02], female:5.14[89%CI 4.09-6.16]), *Bhlhb4^-/-^* grafts produce 4-7 synapses (male:5.49[89%CI 4.17-6.88], female:5.58[89%CI 4.66-6.68]), and *Islet1^-/-^* grafts produce 4-9 synapses (male: 5.78[89%CI 3.81-8.76), female: 5.83[89%CI 4.79-6.69]) per L7^+^ bipolar cell. The probability of synapse varied considerably, perhaps reflective of the stochastic nature of transplantation procedures. On the other hand, estimates for the average number of synapses were more stable and resulted in narrower estimates. Finally, it is noteworthy, that some rod bipolar cells were able to make up to 20 synapses. This value is similar to the number of synapses for a single bipolar cell in the wild type retina, with reported values varying from 25^21^ to 35^22^. Although such robust *de novo* synaptic formation was rare, it highlights the potential for extensive synaptic formation of transplanted grafts.

### De novo host-graft connection remains steady while intra-graft connection decays

Transplanted retinas were examined by MEA recording at 8 and 12 weeks after transplantation in *rd1* or *rd1*;*L7*-GFP mice that were transplanted at 8-12 weeks old. Although no obvious mERG a-waves were detected with the 10 ms pulse stimulation, we consistently observed mERG b-waves as upward peaks of field potential near and over the grafted area, sometimes accompanied by RGC responses (Figure 4a; see Materials and Methods for details). The summary of data (circles) and model predictions (violin plots) for the mERG b-wave probability (Figure 4b) and the response probability of all detected RGCs (Figure 4c) at different conditions shows the model is a reasonable description of collected data. The recorded data is multi-dimensional, with observations being influenced by several predictors (parameters), including graft genotype, graft topology, L-AP4 treatment, stimulus strength, time after transplantation, host genotype, tissue damage, and individual mouse biases. The effect of these predictors was estimated using linear regression. In all of our models we use hierarchical Bayesian inference, which takes into account the hierarchy between predictors. The results of the inference are therefore the joint probability of predictors. We also included b-wave amplitude and spontaneous spiking frequency rates as covariates to estimate the probability of RGC response. Our model shows the RGC response probability is anti-correlated to both the b-wave amplitude and the spontaneous spiking frequency rate (Supplemental Figure 7 covariates). Interestingly, KO grafts exhibited similar RGC responsiveness to wt grafts despite their reduced b-wave activity (Figure 4d; the confidence in the difference in the effect of graft genotype on b-wave amplitude were 90% for *wt-Bhlhb4^-/-^*, 99% for *wt-Islet1^-/-^*, and 87% for *Islet1^-/-^-Bhlhb4^-/-^*), possibly reflecting the presence of a greater number of PKCα^+^ rod bipolar cells in wt grafts than in KO grafts (Figure 2b). Channels within or at the edge of the graft-covered areas have both higher b-wave probability and RGC response probability, indicating these responses are graft-driven (Figure 4e and Supplemental Figure 6 graft topology; confidence in the effect of graft topology were 100% for *on-edge, on-off*, and *edge-off* for both the b-wave and RGC response). L-AP4, the agonistic blocker of glutamate receptor mGluR6 known to be specifically expressed by ON-bipolar cells, largely abolished both b-wave and RGC responses (Figure 4f; confidence in the effect of L-AP4 were 100% for *before-L-AP4* and *L-AP4-after* for both b-wave and RGC response), suggesting the requirement of photoreceptor-bipolar connection for these light responses. The intensity of light stimulus has a clear and significant effect, with stronger stimuli increasing both b-wave and RGC response probability (Figure 4g; confidence in the effect of stimuli were 100% for *weak-strong* for both the RGC response and b-wave) although the effect is less pronounced for the wt graft b-wave (Supplemental Figures 6 and 7 graft genotype and stimulus interaction; confidence for the effect of the interaction is 100% for *wt-Bhlhb4^-/-^* and *wt-Islet1^-/-^* for both *weak* and *strong* stimuli). Intriguingly, the b-wave probability but not RGC responsiveness clearly decreases from 8 weeks post-transplantation to 12 weeks post-transplantation (Figure 4h; confidence for the effect of time after transplantation was 100% for *8W-12W* for the b-wave), possibly indicating that the established host-graft connectivity by 8 weeks post-transplantation remained steady while the intra-graft connections may be lost with time. Unexpectedly, we found a small but noticeable effect on RGC response probability when using the mice with *L7*-GFP labeling as host (Figure 4i; confidence in the effect of *L7*-GFP was 93% for the RGC response), especially when transplanted with wt and *Islet1^-/-^* grafts (Supplemental Figure 7; confidence in the interaction of graft genotype and host genotype were 94% for *wt-Bhlhb4^-/-^* and 99% for *Bhlhb4^-/-^-Islet1^-/-^*). We also confirmed that these RGC responses of transplanted retinas derived from host RGCs by two photon calcium imaging (Supplemental Figure 8).

**Figure 4.**
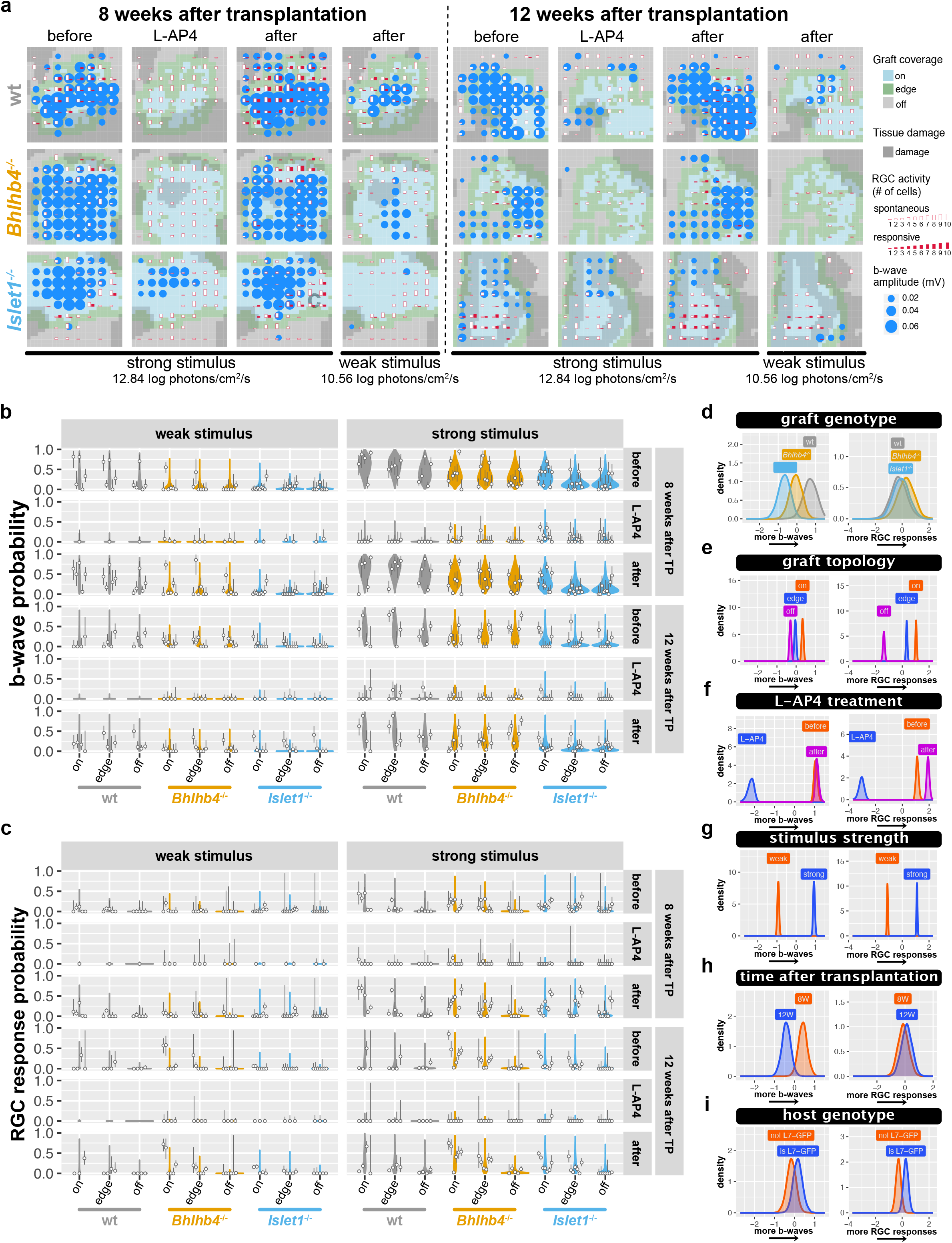
mERG and RGC response to 10 ms pulse stimulus. (a) Examples of MEA recording with graft location, mERG b-wave like response, and RGC response overlaid for each graft genotype. Color tiles represent the graft location (on, edge, off). mERG b-wave like activity by amplitude (> 3-sd) and RGC activity by cell number are shown as circles and bars respectively over the 8×8 electrode locations. Red bars indicate the number of light responsive cells and white bars show the number of cells with spontaneous spiking >3Hz. See Supplemental Figure 5 for more examples of recordings. (b-c) Summary of collected data and the predicted values. Circles indicate the observed probability with bars showing the 89% compatibility interval of binomial trials as calculated by the exact method (Clopper and Pearson). The predictions from our model are shown as violin plots, with wider regions indicating more likely values. (b) shows the probability of observing a b-wave peak on a channel, and (c) shows the probability of RGCs to respond to light. (d-i) Posterior distribution (main effect) of mERG b-wave and RGC response models with 89% compatibility intervals. Full description of the model parameters, including covariates and interaction terms is provided in Supplemental Figures 6 (b-wave model) and 7 (RGC response model). Horizontal axes are in log odds with higher values indicating higher probability of response. n=30167 observations, from 6827 cells, collected from 38 animals (9 wt, 13, *Bhlhb4^-/-^*, and 16 *Islet1^-/-^*) mice (male only), the same mice cohort from the RGC response analysis (Figures 5 and 6).

### ON-pathway activity dominates the host RGC responses in transplanted retinas

Figure 5a shows representative raster plots and peristimulus time histograms of recordings from a normal mouse, an *rd* mouse transplanted with wt and KO grafts, and an age-matched *rd* mouse without transplantation. It is evident that grafts can restore some of the light responsiveness lost in the end-stage *rd* mice. While recordings were carried out with 9-*cis*-retinal for a more stable response, the light response was clearly present even without supplemental 9-*cis*-retinal, indicating that graft photoreceptors can potentially function *in vivo* without direct RPE contact in spite of the rosette formation. We further characterized RGC activity (both responsive and spontaneous) taking into account graft genotype (wt, *Bhlhb4^-/-^*, and *Islet1^-/-^*), L-AP4 treatment, stimulus intensity, and graft coverage. We examined the response pattern (ON, OFF, etc.) of individual cells to the 1s light stimulus through the full before-during-after L-AP4 treatment procedure, and separated RGCs to 4 functional types: not responding to light (*unresponsive*) or being responsive with robust ON responses from the beginning (*on*) or after L-AP4 (*adapted on*), or with response containing the OFF-pathway component (*onXoff*) (see Materials and Methods for detailed description and categorization). Surprisingly, we found the majority of responding RGCs were *on* or *adapted on* types (Figure 5b), with the latter showing ON responses enhanced or emerging after L-AP4 treatment, suggesting the potential of newly formed host-graft synapses to be sensitized. These *adapted on* responses seemed to be more common in *Islet1^-/-^* than in the other grafts, although we cannot be very confident of these differences due to the wide confidence intervals (Supplemental Figure 10 graft genotype; confidence in the effect of graft genotype for *adapted on* response probability was 77% for *wt-Islet1^-/-^* and 83% for *Bhlhb4^-/-^-Islet1^-/-^*). The *onXoff* RGCs, covering only a small portion of responsive RGCs, are suggested to have inputs from the OFF pathways as they responded to the light offset. However, these OFF responses were largely diminished by L-AP4 blockade, implying their dependence of ON-pathway activities, potentially via cross inhibition. Such ON-OFF crossover activities are more often seen in the *Islet1^-/-^* and less frequently in *Bhlhb4^-/-^* lines (Supplemental Figure 10 graft genotype; confidence in the effect of graft genotype for *onXoff* response probability was 90% for *wt-Bhlhb4^-/-^* and *wt-Islet1^-/-^*, and 96% for *Bhlhb4^-/-^-Islet1^-/-^*) and are more strongly associated with graft coverage in KO lines (*Bhlhb4^-/-^* > *Islet1^-/-^* > wt, Supplemental Figure 10 graft genotype and topology interaction).

**Figure 5.**
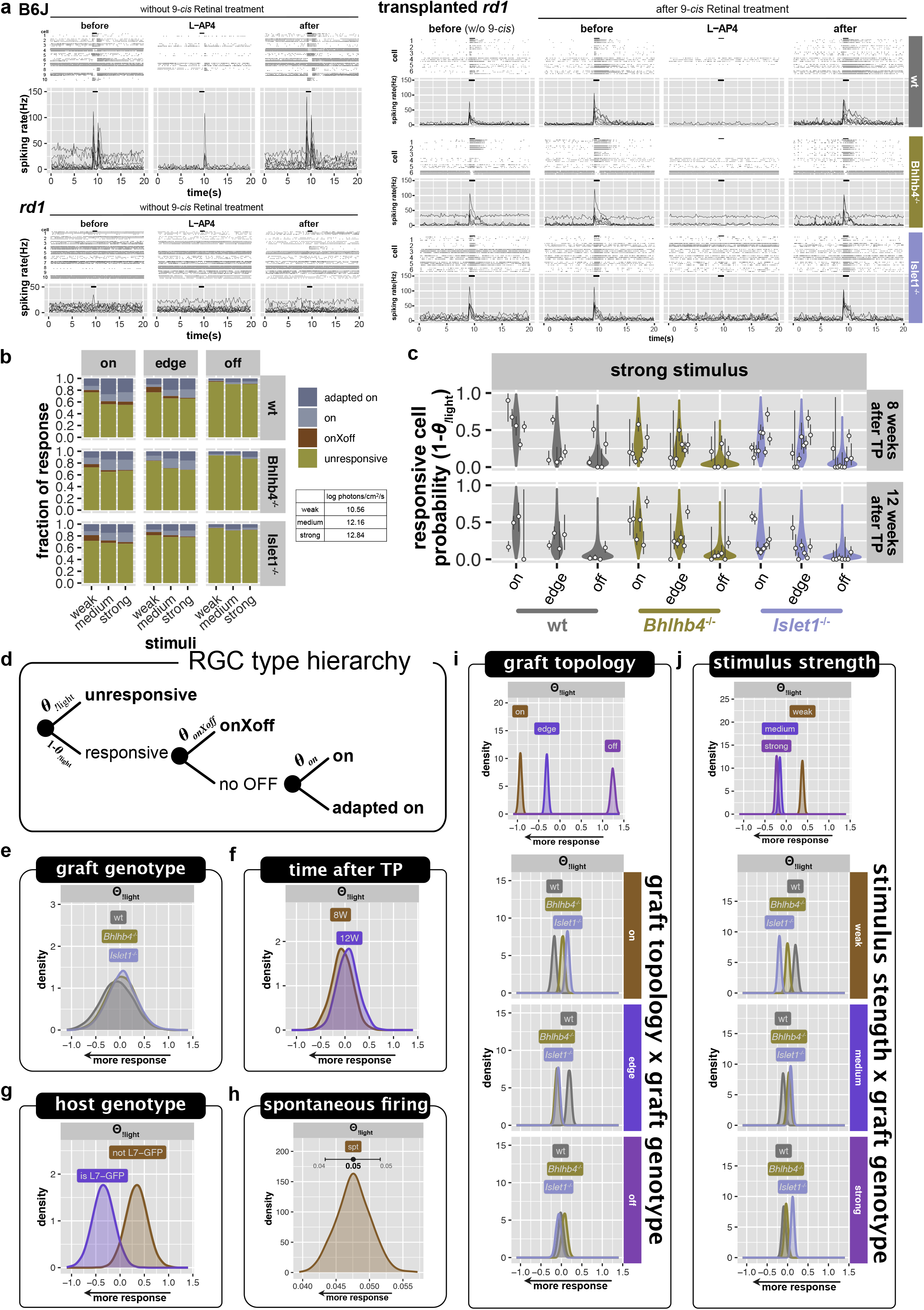
RGC response to 1 s continuous light stimulus. (a) Representative raster plots and peristimulus time histograms of normal mouse, *rd* mouse, and *rd* mouse transplanted with retinal organoids. Raster plots show three repeat recordings from some representative cells. Histogram lines show the spiking frequency of each cell, with bars on top indicating the 1 s light stimulation (strong stimulus for normal and *rd*, and medium stimulus for transplanted mice). (b) Summary of response type ratios of different graft types under different light stimulation and graft coverage from all the recorded samples. (c) Response probability (1 – *θ*_!*light*_) of collected data for strong light stimulation is shown along with model predictions. Circles indicate the observed probability with bars showing the 89% compatibility interval of binomial trials as calculated by the exact method (Clopper and Pearson). The predictions from our model are shown as violin plots, with wider regions indicating more likely values. (d) Hierarchy of RGC response categories for light response pattern. (e-i) Posterior distributions of the parameters for the “no light response” probability (*θ*_!*light*_) for RGC types. For the posterior distributions of the full model parameters including all the response categories, refer to Supplemental Figure 10. Horizontal axes, compatibility intervals, and modes are in log odds. n=16353 observations, from 5738 cells, collected from 38 mice (9 wt, 13, *Bhlhb4^-/-^*, and 16 *Islet1^-/-^*).

### Host RGC responses are strongly associated with graft

Figure 5c shows the summary of collected data and model predictions for the probability of individual RGCs to respond to light. We estimated the effect of predictors using hierarchical Bayesian linear regression, taking into account the dependence between predictors. We used a conditional logit model to estimate the probability of the *unresponsive* and three responsive types (*on*, *adapted on*, and *onXoff*). In this model a simple hierarchical structure among these RGC functional types is assumed as shown in Figure 5d. First, the probability of *unresponsive* RGC (*θ*_!*light*_) is estimated (the exclamation symbol in “!light” denotes the logical NOT operator). Note that 1 – *θ*_!*light*_ is the probability of cells to respond to light and therefore lower values of *θ*_!*light*_ indicate higher responsiveness. Of the cells that respond to light (excluding the *unresponsive* RGCs), cells responding to light offset have a probability *θ_onXoff_* to be *onXoff* type, and the rest of responsive cells (responsive cells excluding *onXoff* type) have either a probability *θ_on_* to be “on” or 1 – *θ_on_* to be *adapted on* type. Figures 5e-j show the posterior distribution of parameters affecting the probability of *unresponsive* RGC (*θ*_!*light*_). Consistent with RGC response to 10 ms stimuli we did not find significant differences between graft genotypes for RGC responsiveness (Figure 5e). There is a clear correlation with graft coverage for response probability (Figure 5i; confidence in the effect of graft topology on the probability of being unresponsive was 100% for *on-edge, on-off*, and *edge-off*). We estimate that upon weak light stimulation cells have approximately 0.03 [89%CI 0.01-0.16] response probability on areas not covered by the graft (off) whereas cells below the graft (on) have a 0.27 [89%CI 0.06-0.62] response probability. In addition to the overall effects of the predictors (the main effects), we further characterize the effect of different graft types by estimating interaction terms with graft genotype. The interaction terms are effects additional to the main effect, under particular combination of predictors. Consequently, although there is a strong overall preference for responses on aeras under and near the graft (on > edge > off, Figure 5i top), wt grafts were characterized by more responses on the “on” graft areas compare to the other grafts, whereas KO grafts show more responses on “edge” graft areas compared to wt grafts (Figure 5i genotype x topology; confidence in the effect of interaction of genotype and topology is higher than 99% for *wt-Bhlhb4^-/-^* and *wt-Islet1^-/-^* for both on and edge locations). So, although the overall responsiveness is similar among graft types, KO lines responses are more dispersed and far-reaching. These light responses were intensity-dependent, with stronger stimuli eliciting more responses (Figure 5j; confidence in the effect of stimulus strength was 100% for *weak-medium* and *weak-strong*, and 93% for *medium-strong*). Interestingly, KO graft-transplanted retinas seem to be more sensitive to low-intensity light (Supplemental Figure 10; confidence in the effect of interaction between graft genotype and stimulus strength was 98% for *wt-Bhlhb4^-/-^* and 100% for *wt-Islet1^-/-^* and *Bhlhb4^-/-^-Islet1^-/-^* for the weak stimulus), whereas the wt had more robust responses under stronger light stimulation. RGC responses remained largely identical at 8 weeks and 12 weeks post transplantation (Figure 5f). Notably, cells with higher spontaneous activity tend to respond less to light (Figure 5h). Lastly, there was an unexpected effect of host genotype, wherein L7-GFP hosts seem to respond better than non-L7-GFP hosts (Figure 5g; confidence in the effect of host genotype on responsiveness was 94%).

### Inputs from Bhlhb4^-/-^ grafts drive host RGC ON responses with better signal-to-noise ratio

Since the increased spontaneous activity in degenerating retina may interfere with light responses, we also analyzed the spontaneous activity of RGCs using the spiking rate of cells from the 9 s recording before light stimulation. Figures 6a and 6b show an example from a sample, illustrating the effect of lowered spontaneous activity by grafts. Figure 6c shows population averages for the three graft types. As the distribution of spontaneous firing follows a lognormal distribution, we estimated the effect of different parameters on the logmean. Figure 6d shows the distribution of RGCs firing frequency for responsive and unresponsive cells for different graft lines, overlaid with model predictions (see Supplemental Figure 11 for the same plot in log frequency). In this analysis we used light responsiveness instead of graft topology, as light responsiveness is a more accurate indicator that RGCs have connections that lead to graft photoreceptors. Our analysis indicates that higher spontaneous activity is reduced in host retina transplanted with *Bhlhb4^-/-^* (Figure 6e, confidence in the effect of graft genotype was 87% for *wt-Bhlhb4^-/-^* and 92% for *Bhlhb4^-/-^-Iselet1^-/-^*). Spontaneous activity clearly decreased after L-AP4 washout (Figure 6g, confidence in the effect of L-AP4 were 100% for *before-after* and *L-AP4-after)*, and we speculate that this is related to the sensitization of host-graft synapses by L-AP4, followed by a dis-inhibitive potentiation of bipolar cells after washout. Similar to RGC responses, spontaneous firing remained largely identical at 8 weeks and 12 weeks after transplantation (Supplemental Figure 11 time after TP). There is a clear and substantial difference between cells that respond to light (*on*, *adapted on*, *onXoff*) and cells that do not (*unresponsive*), with responsive cells having lower spontaneous activity (Figure 6e and Supplemental Figure 11 responsiveness; confidence in the effect of light responsiveness was 100%). Interestingly, there was a small but distinctive effect of graft genotype on the effect of light responsiveness, with KO grafts having lower spontaneous firing rates in unresponsive cells (Figure 6e and Supplemental Figure 11 graft genotype responsive interaction; confidence in the effect of graft genotype was higher than 99% for *wt-Bhlhb4^-/-^*, *wt-Islet1^-/-^*, and *Bhlhb4^-/-^-Islet1^-/-^* for both responsive and unresponsive cells).

**Figure 6.**
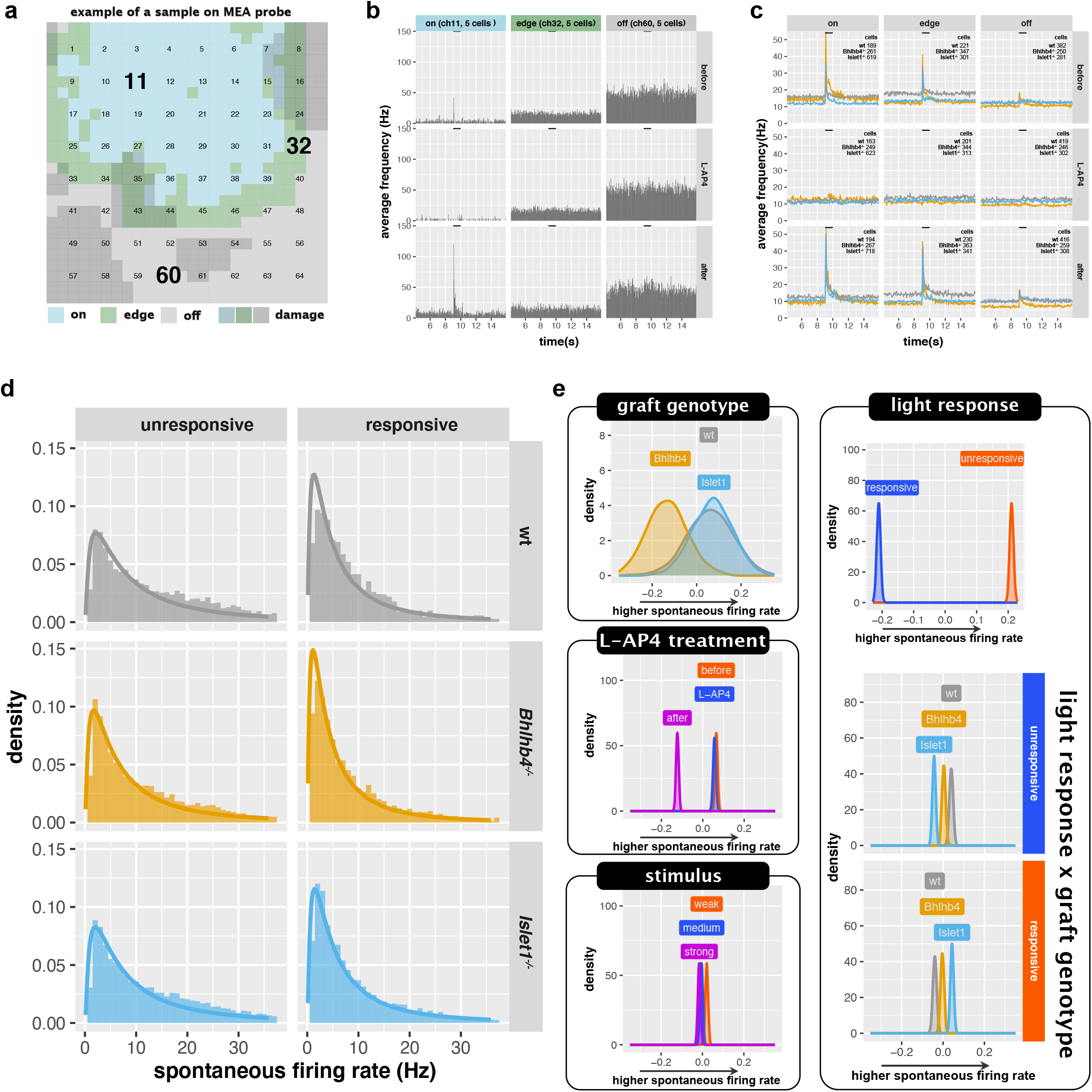
RGC spontaneous firing before light stimulation. (a-b) Examples of MEA recordings. (a) shows the position of graft (on, edge, off) on the 64 electrodes of the MEA probe. The dark gray areas indicate areas where tissue damage was apparent (see Materials and Methods for details). (b) Shows the peristimulus histogram of three channels (11, 32, and 60, each with 5 cells detected) which correspond to on, edge, and off graft areas respectively. (c) shows population averages of spontaneous firing rate (the height of the baseline) and light responses (peaks triggered by onset and offset of light stimulus) on different graft areas (on, edge, off) and before, during, and after L-AP4 treatment. The bar on top indicates the 1 s light stimulation. This is the average of responses to samples collected 12 weeks after transplantation and stimulated with the strong light condition. See Supplemental Figure 9 for population averages of other conditions. The spontaneous activity (defined as the average firing frequency prior to light stimulation) was modeled with a lognormal distribution. (d) shows the histogram of the spontaneous firing rate overlayed with the modeled lognormal distribution for responsive and unresponsive of host transplanted with different graft lines. The same plot but in with the horizontal axis in log is shown in Supplemental Figure 10. (e) Posterior distributions of the parameters affecting the logmean. For the posterior distributions of the full model parameters refer to Supplemental Figure 11. n=44213 observations, from 5738 cells, collected from 38 mice (9 wt, 13, *Bhlhb4^-/-^*, and 16 *Islet1^-/-^*).

### Mice transplanted with Bhlhb4^-/-^ and Islet1^-/-^ grafts perform better at light avoidance response

While KO grafts did not show a noticeable improvement on the RGC light response over wt grafts, they seemed to better suppress high frequency spontaneous spiking. We speculated that KO graft transplantation would result in better functional recovery due to its reduced noise. To test this, we conducted a behavior examination of light avoidance (Figure 7, see Materials and Methods for details). Mice were placed in a cage with two chambers which are randomly illuminated by mesopic light proceeded by a mild electric shock. Mice that move often, indicated by higher inter trial interval (ITI) counts, can avoid the electric shock purely by chance as can be seen in Figure 7a for the negative control. However, mice that can perceive the light learn to avoid the electric shock using the light as a cue, shifting this curve upwards. We modeled the effect of each group (control, wt, *Bhlhb4^-/-^*, and *Islet1^-/-^*) on the success probability while accounting for the ITI and other factors as predictors (Supplemental Figure 12). Consistent with our hypothesis we found that mice transplanted with KO grafts performed better than wt grafts (*Bhlhb4^-/-^* > *Islet1^-/-^* > wt, Figures 7b and c; confidence in the effect of graft/group was 88% for wt-*negative*, 98% for *Islet1^-/-^-negative*, 100% for *Bhlhb4^-/-^-negative*, 80% for *wt-Islet1^-/-^*, 97% for *wt-Bhlhb4^-/-^*, and 81% for *Bhlhb4^-/-^-Islet1^-/-^*). We found that females performed somewhat better than males (Supplemental Figure 12; confidence in the effect of sex was 89%). In females, both *Bhlhb4^-/-^* and *Islet1^-/-^* grafts show a modest improvement over wt grafts, whereas the improvement is only apparent in *Bhlhb4^-/-^* in males (Figure 7d).

**Figure 7.**
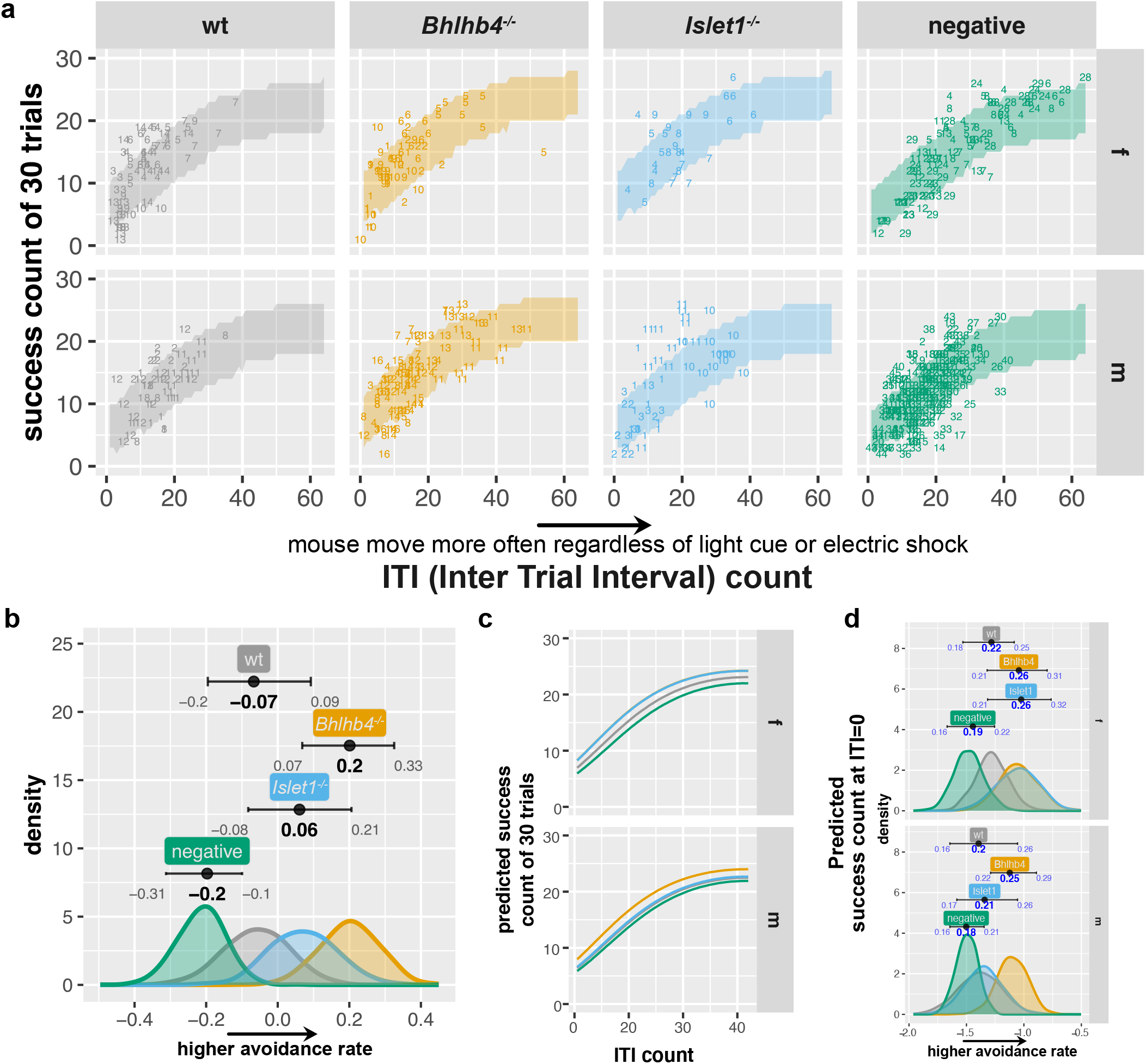
Light Avoidance Test. Mice are placed in dark box and presented with a light cue before a mild electric shock. Mice that can sense the light learn to avoid the electric shock using the light cue. (a) The number of successful avoidances out of 30 trials is plotted against inter trial interval (ITI), which represent the sporadic movements of the mouse. Numbers indicate the id of a mouse for each group. Success count increases with ITI as mice that move often may avoid the shock purely by chance. Mouse that perceive light and learn to use it as a warning can increase their success count with lower ITI. The number of success counts was modeled using the binomial distribution using group, sex, mouse, ITI, and trail number as predictors, and the shaded area shows the 89% predicted success count. (b) Estimated effect of each group (*β_grp_*) with 89% compatibility intervals. Distribution for the rest of the parameters is provided in Supplemental Figure 12. Note the horizontal axis is the log odds. (c) Overlay of predicted mean avoidance. (d) Predicted success rate, at ITI=0, for male and female mice. Note that the bottom axis is logit (log odds). The compatibility interval value and modes (in blue) were transformed to probability [0-1] (*logistic*(*β*_0_ + *β_grp_* + *β_sex_* + *β_grpXsex_*)) to facilitate interpretation. The number of mice used for this analysis was 86 mice (14 wt, 15 *Bhlhb4^-/-^*, 11 *Islet1^-/-^*, and 45 age-matched negative control (*rd1*)).

## Discussion

In this study we transplanted to end-stage degenerated retinas, minimizing the likelihood of observing the effects of remaining host photoreceptor cells or cytoplasmic material transfer rather than neural integration of grafted cells^23–26^. The use of end-stage *rd1*; *L7*-GFP model conclusively and unequivocally proves the neural integration of grafted cells, which is further substantiated by electrophysiology and behavior experiments. We prepared *Bhlhb4^-/-^*, and *Islet1^-/-^* grafts aiming to reduce the population of bipolar cells to improve host-graft contact. The PKCα^+^ rod bipolar cell population was markedly reduced as expected, although photoreceptor density remained the same. It is possible that nascent progenitor cells are diverted to alternative cell fates or remain partially differentiated, unable to commit to a final cell fate in *Bhlhb4^-/-^* and *Islet1^-/-^* grafts^18,27^ Overall, however, *Bhlhb4^-/-^* and *Islet1^-/-^* grafts displayed normal photoreceptor differentiation and retinal structure.

KO grafts did not substantially increase the number of host bipolar cells forming synapses, or improved RGC light responsiveness. Instead we found that KO grafts allow higher number of synapses per host bipolar cells, possibly due to the higher number of graft photoreceptors without intra-graft partners. The higher number of synapses per rod bipolar is also consistent with the higher light-responsiveness to weak light stimuli observed in KO-graft transplanted retinas. On the other hand, KO graft responses were characterized by reduced spontaneous activity. Degenerated retinas are characterized by increased spontaneous activity owing to the lack of photoreceptor input^28–30^. It is tempting to speculate that bipolar cells were able to establish more connections in grafts deprived of bipolar cells, resulting in sufficient photoreceptor input to restore inner retinal signaling and to reduce aberrant RGC activity. This in turn may be reflected in the better performance of KO grafts transplanted animals in behavior tests. The reduction in spontaneous activity (Figure 6) and better performance in behavior test (Figure 7) seem to indicate *Bhlhb4^-/-^* is more advantageous than *Islet1^-/-^*, at least in male mice, although synaptic connectivity was similar. It may be that we were unable to observe differences in connectivity due to the large sample variation, or alternatively it may be that although the number of synapses is similar, the quality, i.e. the strength of synaptic connectivity^20^ or the distribution of synapse across rosettes differs. Although we are uncertain of the underlying mechanism, reduction in aberrant RGC activity seems to substantially improve the performance after transplantation. This highlights the importance of future approaches to focus on the reconstruction of the inner retina circuitry in addition to neural integration of transplanted cells, as degenerate retinas undergo neural remodeling^31,32^. Our results indicate the possibility that remodeled inner retinal circuitry can still be reengaged with sufficient photoreceptor input.

KO grafts display reduced b-wave activity despite the similar RGC light responsiveness, consistent with the reduction in bipolar cell population in the graft (wt>*Bhlhb4^-/-^*>*Islet1^-/-^* for both b-wave and bipolar cells; see Figure 2 and Figure 4). We think the difference between b-wave and RGC responsiveness is reflective of intra graft photoreceptor-bipolar cell connections. These intra graft connections not only deprive the host retina from forming contacts with graft photoreceptor cells but may also have some adverse effects on the host retinal activities. Our analysis of the spontaneous activity shows that RGC cells that are light responsive with synaptic inputs originating from graft photoreceptors, have a clear reduction in spontaneous activity. However, in unresponsive host cells without effective inputs from grafts, the spontaneous activity is higher wherein more presumptive intra graft photoreceptor-bipolar connections exist. Although we are not certain of the mechanism that leads to more spontaneous activity in grafts with more intra graft connections, these results highlight the benefits of KO grafts.

During the course of our experiments, we found unexpected effects of host genetic background. Firstly, host sex seems to affect synapse formation and possibly the behavior response, with females performing better than males. Secondly, *rd1*;*L7*-GFP mice seem to perform better in MEA recordings than *rd1* mice although these mice have the same C57BL/6J background. It is possible that the *L7*-GFP may retain some FVB/N background. We do not currently understand how these differences originate but it underscores the need to control for these variables.

The requirement of establishing synaptic integration is a significant challenge for photoreceptor cell therapies. We have re-enforced our previous report, showing conclusively that stem cell derived retinas form synaptic connections with the host to restore light responsiveness^12^. Furthermore, we have expanded our analysis and shown that aberrant firing is reduced by grafts. This effect is further enhanced in genetically engineered cell lines, perhaps due to improved input to bipolar cells and therefore improved outcome of transplantation. Our genetically engineered cell lines are unlikely to result in deleterious effects, as they are not introducing an exogenous gene, but rather deleting part of a gene that is expressed in a subset of cells in the graft. Furthermore, in the reported *Bhlhb4^-/-^* or *Islet1^-/-^* mice, there are no known unfavorable ocular abnormalities other than the reduction or absence of the b-wave, and both morphology and function of photoreceptors seem normal. Local transplantation of these tissues is therefore well justified considering the potential benefits in patients at the risk of becoming blind.

## Acknowledgments

We thank Dr. Genshiro A Sunagawa and Prof. Chuan-Chin Chiao for critical reading and comments on the manuscript. This work was supported by JSPS KAKENHI Grant Number 15K10913, JK19K09942, and 17K16994.

## Author Contributions

MM and MT conceived the study. HYT and TM performed electrophysiological experiments. TH and JSun prepared 3D retinas. MM and JSun designed, constructed cell lines and transplanted 3D retinas. AO procured the *Thy1*-GCaMP3 mouse line. JSho maintained animals and assisted on transplantation. TH and MF performed RT-PCR. TM conducted statistical analysis. TH, RA, MM, JSun and HYT performed immunostaining, imaging and quantification. TM, MM and HYT wrote the manuscript

## Declaration of Interests

There is potential Competing Interest.

The corresponding author is currently filing for a patent regarding the genetically modified retinal organoids.

Information of Patent applicant applicant: RIKEN, SUMITOMO DAINIPPON PHARMA

CO., LTD name of inventors: Michiko MANDAI, Masayo TAKAHASHI, Suguru

YAMASAKI

application number: PCT/JP2017/042238

status of application: national phase

specific aspect of manuscript covered in patent application: The phenotype of retinal grafts with reduced inner cell population and the probability of improved results following retinal transplantation are covered by the above patent.

---

**Supplemental Figure 1.**
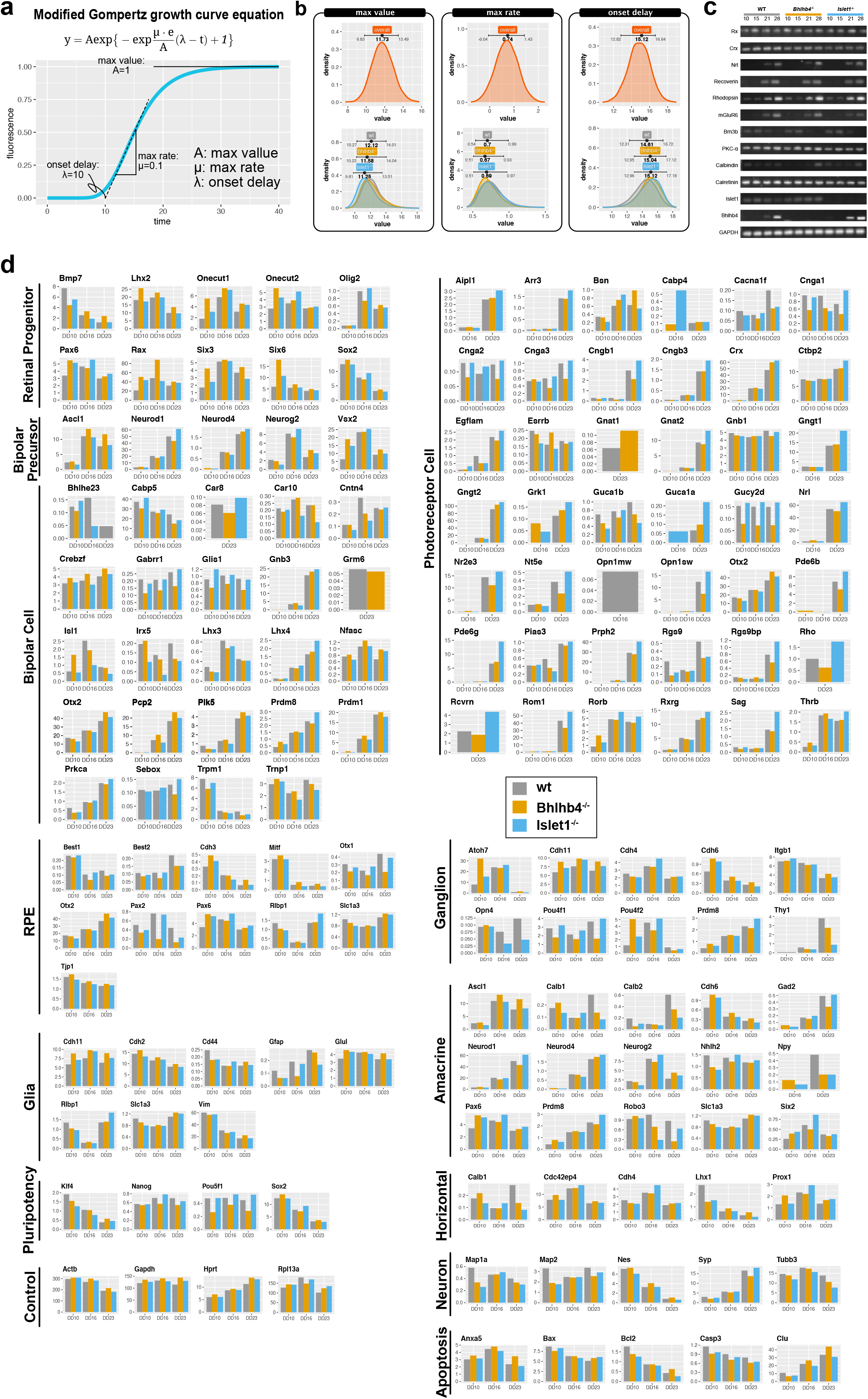
Characterization of retinal organoids. (a) Growth curve model of GFP signal from retinal organoids. The modified Gompertz equation is defined by 3 parameters: *A* the max value, *μ* the max rate, and *λ* the onset delay. *Nrl*-GFP signal increase was modeled using this growth curve assuming a Gamma distribution. (b) Posterior estimates for parameters of growth curve. Upper graphs show the overall estimated value and the lower graph shows the estimate for different cell lines. Bars indicate the 89% compatibility interval and the point is the mode. n=170 organoids (54 wt, 61 *Bhlhb4^-/-^*, and 55 *Islet1^-/-^*) were used for this analysis. (c) RT-PCR of key genes from organoids differentiated from a clone of ROSA26^+/*Nrl*-CtBP2:tdTomato^ ES cells. (d) Bar charts showing expression pattern of wt, *Bhlhb4^-/-^*, and *Islet1^-/-^* retinal organoids at DD10, DD16, and DD23 from microarray analysis. Bars indicate the relative expression of genes using the 75^th^ percentile normalization.

**Supplemental Figure 2.**
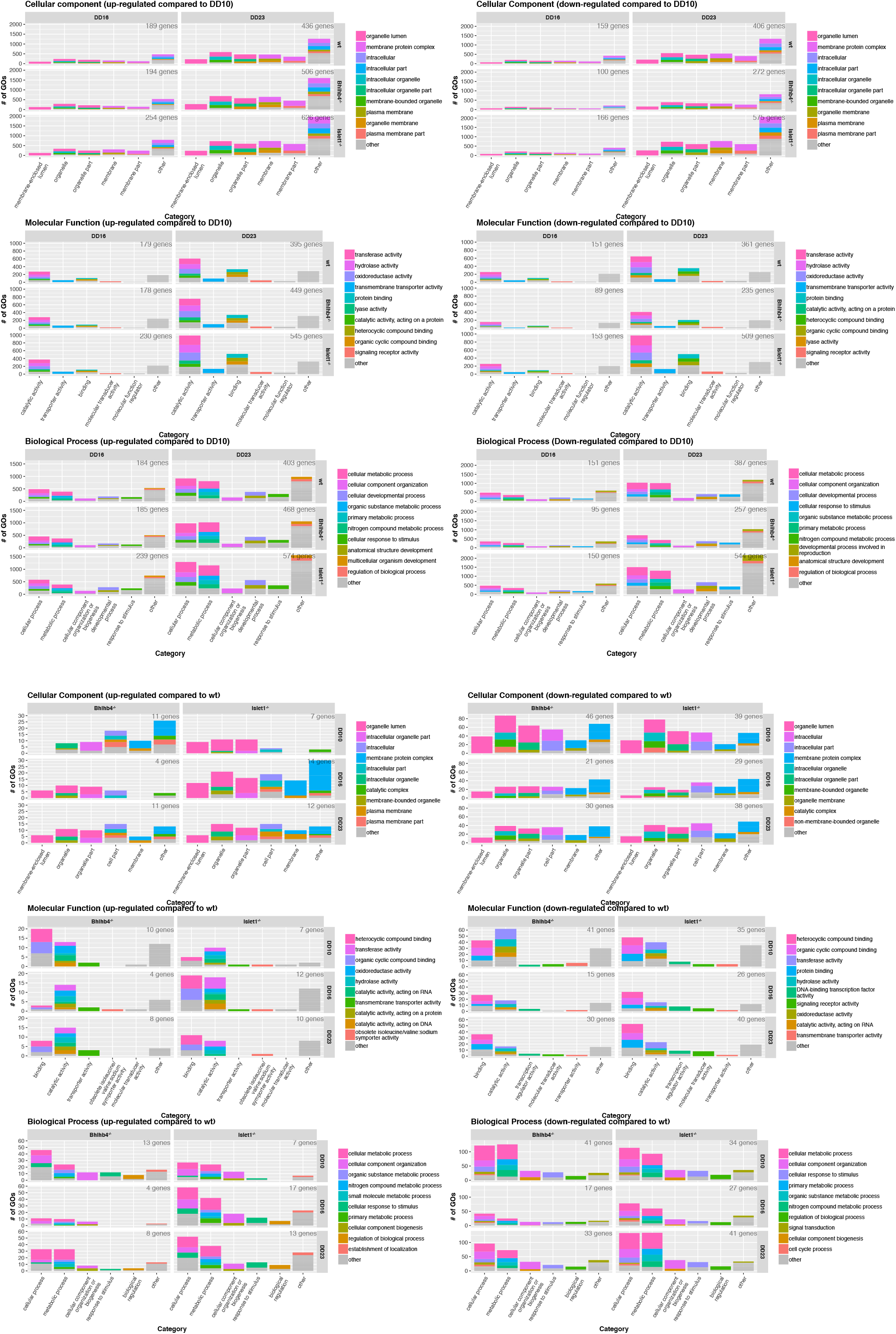
GO enrichment analysis of Molecular function (MF), Cellular component (CC), and Biological process (BP) of differentially expressed genes with more than 4-fold change (points outside the lines in Figures 1e and 1f). Graphs on the left show up-regulated genes and graphs on the right show up-regulated genes. Top 3 rows show the expression change of DD16 and DD23 compared to DD10 (Figure 1e), whereas the bottom 3 rows show the expression of *Bhlhb4^-/-^* and *Islet1^-/-^* compared wt (Figure 1f).

**Supplemental Figure 3.**
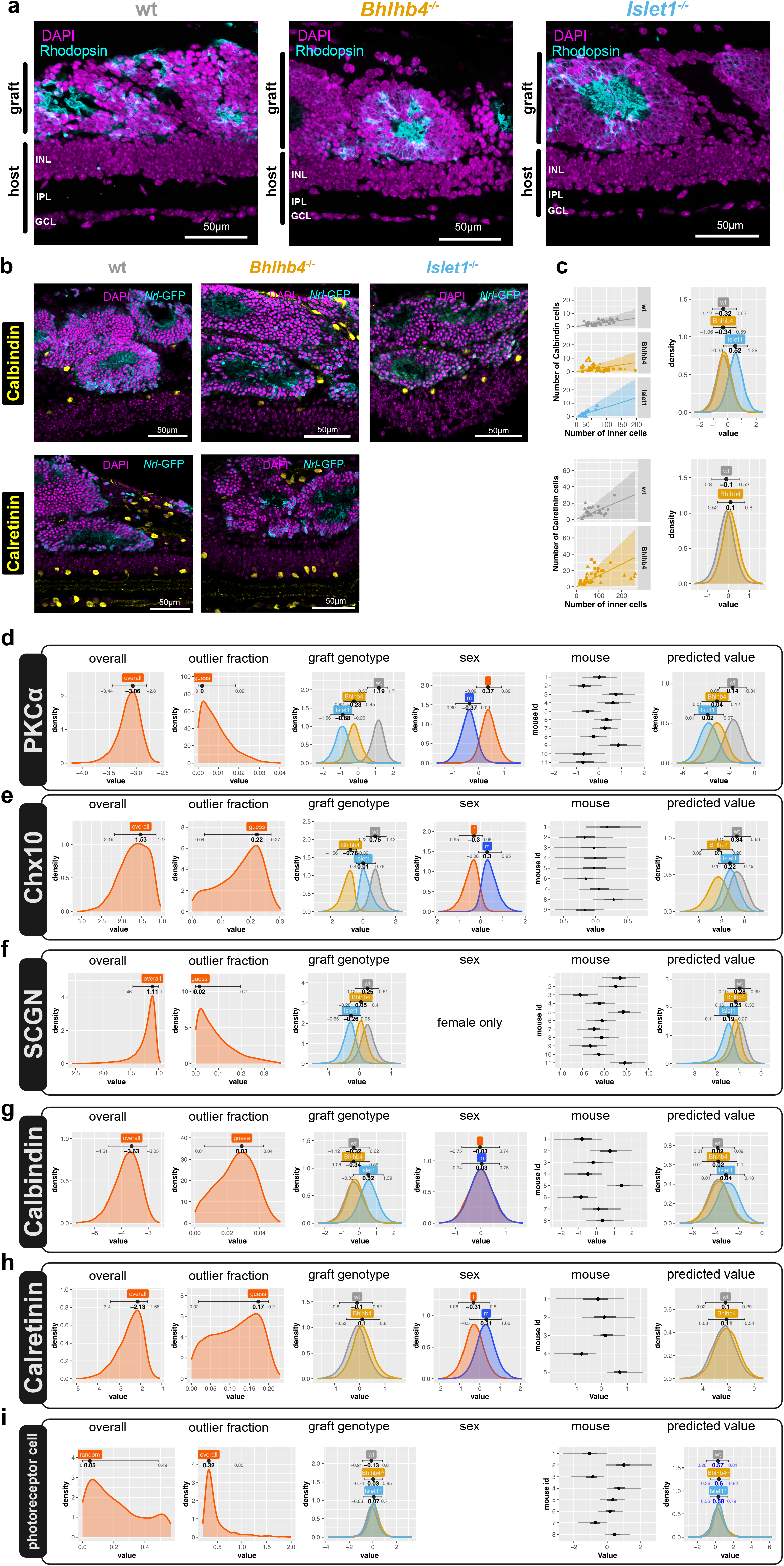
Characterization of graft cells after subretinal transplantation. (a) Representative images for transplanted grafts forming rosette like structures with photoreceptor cells showing polarized rhodopsin expression at inner and outer segment (IS/OS) like structures towards the center of rosettes. (b) Calbindin^+^ and Calretinin^+^ staining in wt, *Bhlhb4^-/-^*, and *Islet1^-/-^* grafts. (c) Calbindin^+^ cells (top) and Calretinin^+^ cells (bottom) were plotted against the number of inner cells in the graft. Dots indicate the count value for a given image (section). The line represents the most likely value and the shaded area represents the 89% compatibility interval assuming that the number of positive cells for a positive marker follows the binomial distribution given the number of total (inner) cells. Right plots show the estimated effect of graft genotype on the probability of the marker being positive, with mode and 89% compatibility intervals indicated on top. Note that the horizontal scale (value) is logit, i.e. log odds, with higher values indicating higher probability. n=102 cells from 8 mice (3 wt, 3 *Bhlhb4^-/-^*, and 2 *Islet1^-/-^*) for Calbindin^+^ cells and n=68 cells from 5 mice (2 wt and 3 *Bhlhb4^-/-^*) for Calretinin^+^ cells. (d-h) show the model parameters and posterior estimates for the analysis of PKCα^+^, Chx10^+^, SCGN+, Calbindin^+^, and Calretinin^+^ cells, and (i) shows the parameters for a similar model for the number of photoreceptor cells in the graft (photoreceptor cells vs total graft cells). The number of inner cells for a particular marker was analyzed assuming a linear relationship between the number of graft cells and the number of cells positive for the particular marker using the binomial distribution as likelihood. Graphs show the posterior distribution for the model parameters with mode and 89% compatibility interval indicated on top. The horizontal axis for “guess” represents the fraction [0-1] of data points that are estimated to be generated by a random process, i.e. outliers. The horizontal axis for “overall” (*β*_0_), “graft genotype” (*β_lin_*), “sex”(*β_sex_*), “mouse” (*β_mus_*), and “predicted value” are in logit scale (log odds), as the data was modeled using the logistic function as link function. Higher values indicate higher fraction of positive cells. The “predicted value” indicates the expected value for a given line (*β*_0_ + *β_lin_*). The compatibility interval value was transformed to probability [0-1] (*logistic*(*β*_0_ + *β_lin_*))to facilitate interpretation.

**Supplemental Figure 4.**
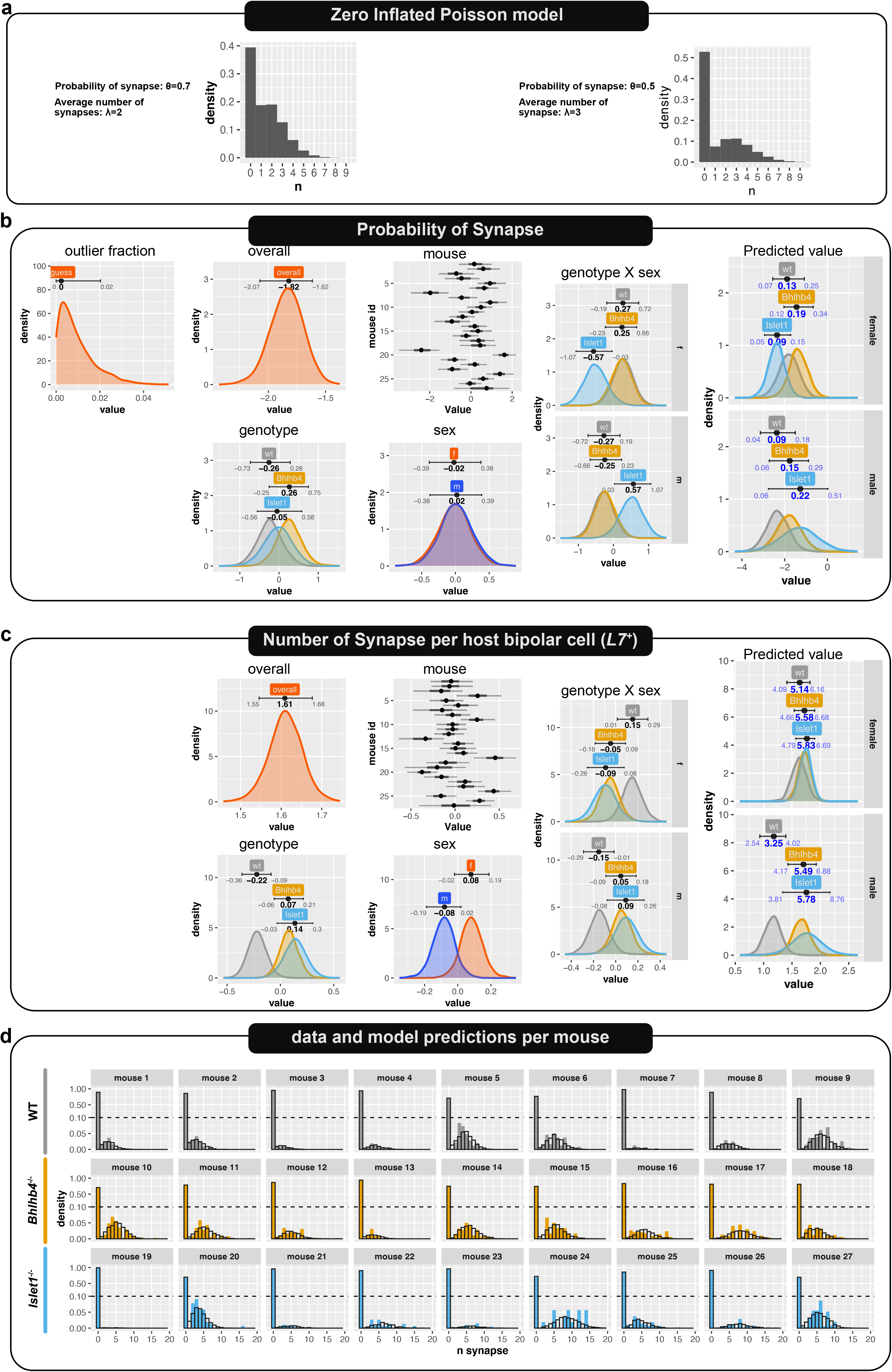
Synapse number per bipolar cell analysis. The number of graft-host synapses was analyzed with a Zero-Inflated Poisson model as the data contain a large number of zero counts. (a) Example of Zero-Inflate Poisson distribution. In this framework, the probability of making a synapse is assumed to be independent of the average number of synapses per bipolar cell. The data is described by the probability of making a synapse *θ* and the average number of synapses *λ*, estimated from the data using “graft genotype” (*β_lin_*), “sex”(*β_sex_*), “mouse” (*β_mus_*) as predictors. Graphs in (b) and (c) show the posterior distribution for the model parameters with 89% compatibility interval and mode indicated on top of each posterior distribution for the probability of synapse *θ* (*b*) and the average number of synapses *λ* (c). The horizontal axis for “guess” represents the fraction [0-1] of data points that are estimated to be generated by a random process, i.e. outliers. The horizontal axis for “overall” (*β*_0_), “graft genotype” (*β_lin_*), “sex”(*β_sex_*), “mouse” (*β_mus_*), “genotype x sex”(*β_linXsex_*), and “predicted value” are in log odds for the probability of synapse and in log for the average number of synapses. “genotype x sex” is the interaction term of genotype and sex, i.e. a secondary conditional effect in addition to the main effect of graft genotype and sex under different combinations of graft genotype and sex. The “predicted value” indicates the expected value of different grafts transplanted to host of different sex (*β*_0_ + *β_lin_* + *β_sex_* + *β_linXsex_*). The confidence interval and mode for the predicted value (in blue) were transformed to their original data scales, showing the estimated probability of synapse (*logistic*(*β*_0_ + *β_lin_* + *β_sex_* + *β_linXsex_*)) and the average number of synapses per host bipolar cells (exp(*β*_0_ + *β_lin_* + *β_sex_* + *β_linXsex_*)). (d) Breakdown of collected data per mouse shows the large variance from sample to sample. Filled bars show the data and bars without fill show the model predictions, indicating the model is reasonably representing the data.

**Supplemental Figure 5.**
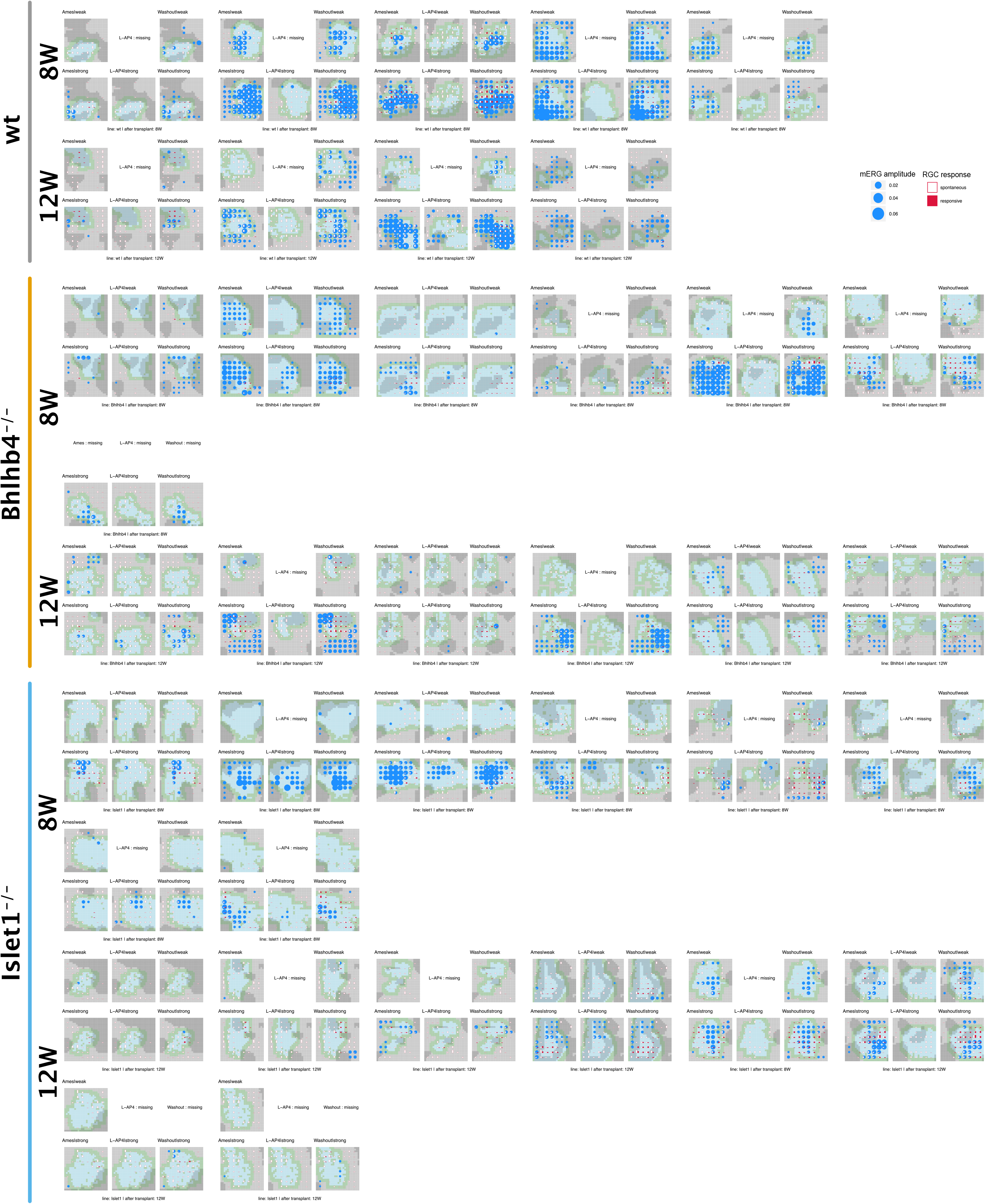
mERG and RGC response to 10 ms stimulus. The graphs show recordings with graft location, mERG b-wave like response, and RGC response overlaid. Color tiles represent the location of the graft on the 8×8 MEA array. Gray: not covered by the graft, light blue: covered by the graft, olive-green: graft edges, dark gray: recorded response may be unreliable due to tissue damage, optic disc, etc. mERG b-wave like activity by amplitude (> 3-fold sd) and RGC activity by cell number are shown as circles (0.02, 0.04, 0.06 mV) and bars respectively over the electrode locations. Bars indicate the number of light responsive cells (red) and the number of cells with spontaneous activity (>3Hz).

**Supplemental Figure 6.**
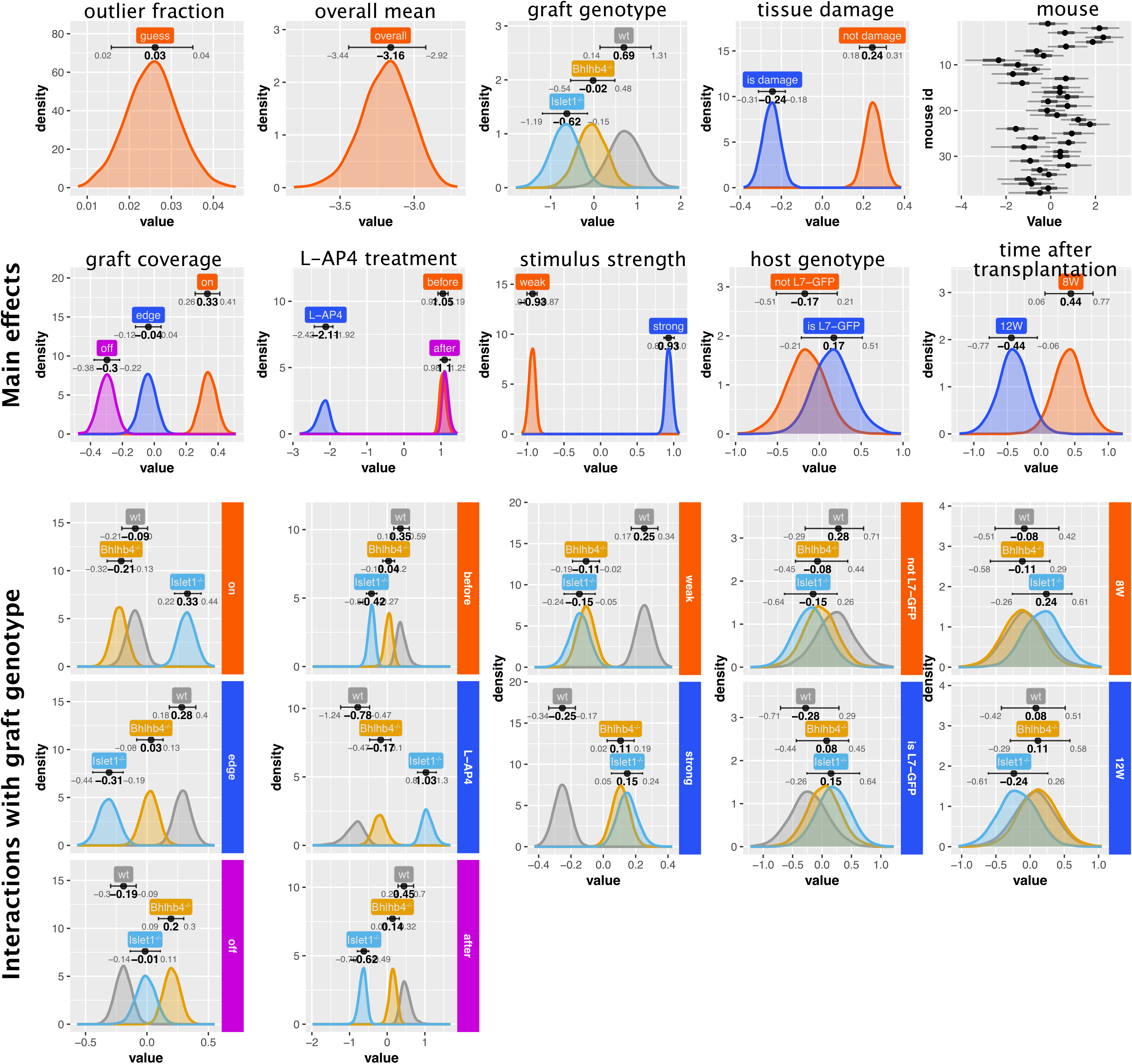
mERG b-wave analysis. The probability of observing a b-wave was modeled with a Bernoulli model using “graft genotype” (*β_lin_*), “tissue damage”(*β_dmg_*), “mouse” (*β_mus_*), “graft coverage” (*β_tpl_*), “L-AP4 treatment”(*β_cnd_*), “stimulus strength”(*β_stm_*), “tissue damage”(*β_dmg_*), “host genotype”(*β_17g_*), “time after transplantation”(*β_tim_*). Graphs show the posterior distribution of estimated parameters with 89% compatibility intervals and mode. The top two rows show the main effects. For graft coverage (on, edge, or off), L-AP4 treatment (before, L-AP4, or after), stimulus strength (weak or strong), host genotype (*L7*-GFP or not), and time after transplantation (8 weeks or 12 weeks) we calculated the interaction terms with graft genotype. These interaction terms with graft genotype are shown under their respective main effects’ plots. Note that interactions are conditional effects that are present under a particular combination of predictors in addition to the main effects. The horizontal axis for “guess” represents the fraction [0-1] of data points that are estimated to be generated by a random process, i.e. outliers. The horizontal axis for “overall” (*β*_0_), “graft genotype” (*β_lin_*), “tissue damage”(*β_dmg_*), “mouse” (*β_mus_*), “graft coverage”(*β_tpl_*), “L-AP4 treatment”(*β_cnd_*), “stimulus strength”(*β_stm_*), “tissue damage”(*β_dmg_*), “host genotype”(*β_17g_*), “time after transplantation”(*β_tim_*), and their interaction terms are in log odds, with larger values indicating higher probabilities.

**Supplemental Figure 7.**
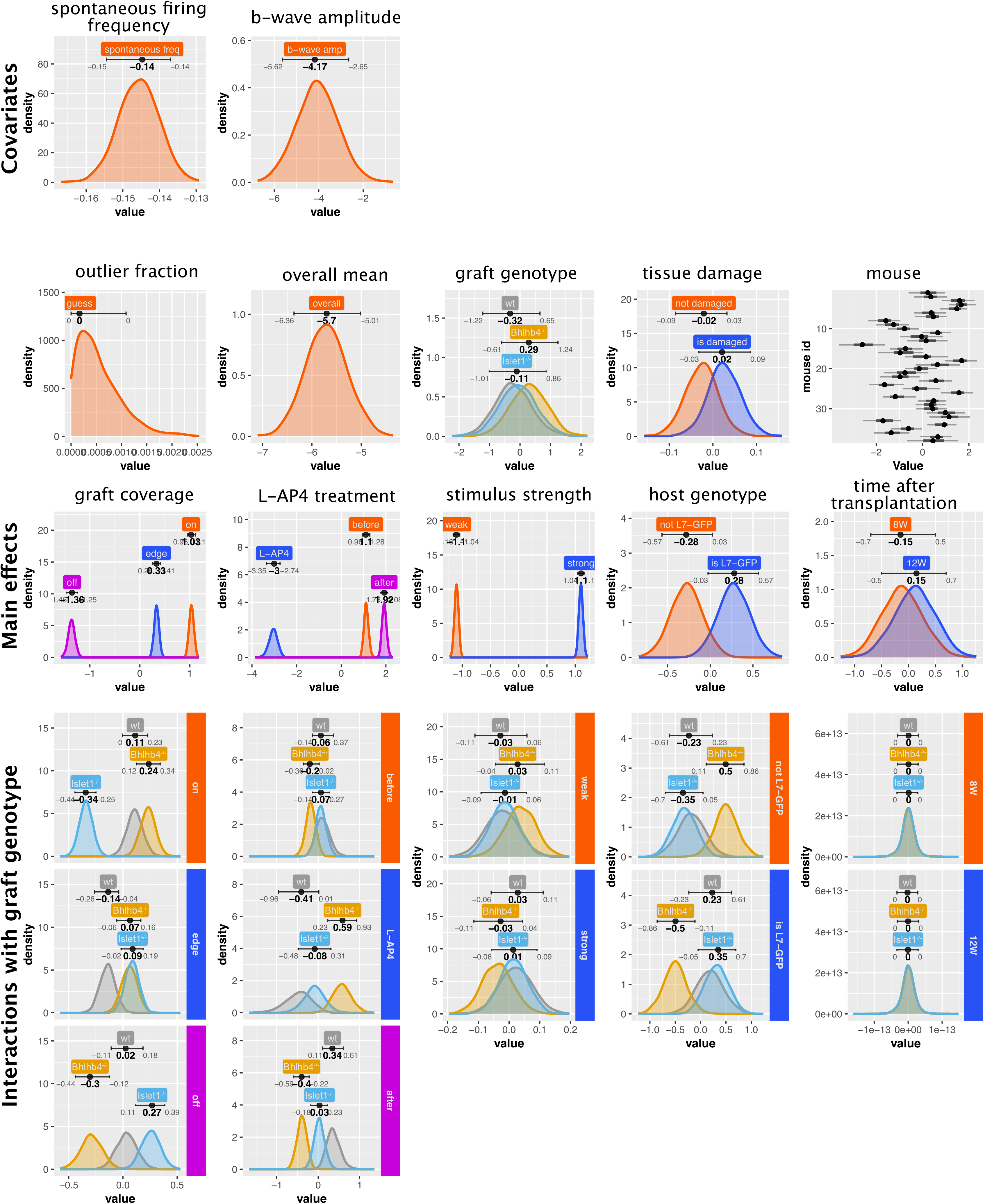
Modeling of RGC response to pulse stimulus (10 ms). The probability of observing an RGC response was modeled with a Bernoulli model, using the “spontaneous firing frequency” (*β_spt_*), “b-wave amplitude” (*β_bwv_*), “graft genotype” (*β_lin_*), “tissue damage”(*β_dmg_*), “mouse” (*β_mus_*), “graft coverage”(*β_tpl_*), “L-AP4 treatment”(*β_cnd_*), “stimulus strength”(*β_stm_*), “tissue damage”(*β_dmg_*), “host genotype”(*β_17g_*), and “time after transplantation”(*β_tim_*) as predictors. Graphs show the posterior distribution of estimated parameters with 89% compatibility interval and mode. The top row shows the covariates (continuous variables) and the second and third rows shows main effect of categorical variables. For graft coverage (on, edge, or off), L-AP4 treatment (before, L-AP4, or after), stimulus strength (weak or strong), host genotype (*L7*-GFP or not), and time after transplantation (8 weeks or 12 weeks) we calculated the interaction terms with graft genotype. These interaction terms with graft genotype are shown under their respective main effects’ plots. Note that interactions are conditional effects that are present under a particular combination of predictors in addition to the main effects. The horizontal axis for “guess” represents the fraction [0-1] of data points that are estimated to be generated by a random process, i.e. outliers. The horizontal axis for “overall” (*β*_0_), “graft genotype” (*β_lin_*), “tissue damage”(*β_dmg_*), “mouse” (*β_mus_*), “graft coverage”(*β_tpl_*), “L-AP4 treatment”(*β_cnd_*), “stimulus strength”(*β_stm_*), “tissue damage”(*β_dmg_*), “host genotype”(*β_17g_*), “time after transplantation”(*β_tim_*), and their interaction terms are in log odds, with higher values indicating higher probabilities. n=30167 observations, from 6827 cells, collected from 38 animals (9 wt, 13, *Bhlhb4^-/-^*, and 16 *Islet1^-/-^*) mice (male only), the same mice cohort from the RGC response analysis (Figures 5 and 6).

**Supplemental Figure 8.**
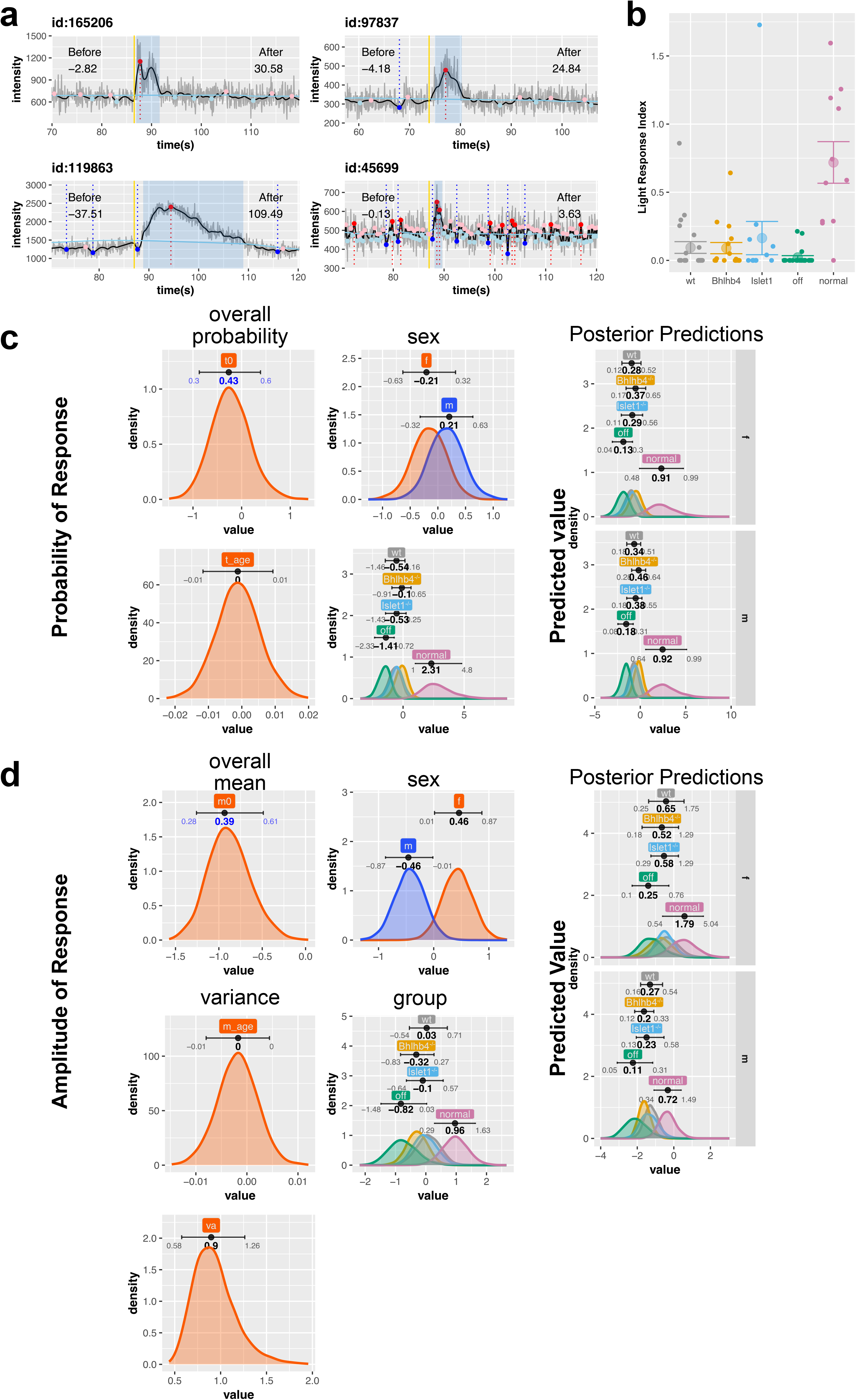
Two photon measurement of RGC calcium change. (a) Example of fluorescence time course in a ROI. The gray trace shows the raw data, the black trace shows the slightly smoothed data, and the light blue trace shows the baseline (strongly smoothed) trace. Pink and light blue points show local maxima and minima, and red and blue points with dotted lines indicate peaks with amplitude exceeding baseline sd. The yellow line indicates the position of the light stimulus, and the blue shaded area shows the detected peaks (maxima exceeding three times the baseline sd following light stimulus). Numbers in the graph correspond to the calculated fluorescence intensity (area) before and after the light stimulus. Note that peak amplitude and kinetics vary substantially. (b) Summary of recorded light responses for samples collected from *rd1;Thy1*-GCaMP3 animals transplanted with wt, *Bhlhb4^-/-^, Islet1^-/-^* grafts, and age-matched control animals (normal, B6J). The data in (b) was modeled with a gamma hurdle model, and (c) shows the posterior distribution of parameters for the probability of a response, and (d) shows the posterior distribution of predictors for the amplitude response, with 89% compatibility intervals and mode indicated on top of the distribution. For the predicted value and overall parameters, the intervals show the values converted to probability (*logistic*(∑*β_predictos_*)) for the probability (b) and amplitude (exp(∑*β_predictors_*)) for the amplitude (a). The number of mice used for this analysis was 63 (22 wt, 17 *Bhlhb4^-/-^*, 13 *Islet1^-/-^*, and 11 negative control).

**Supplemental Figure 9.**
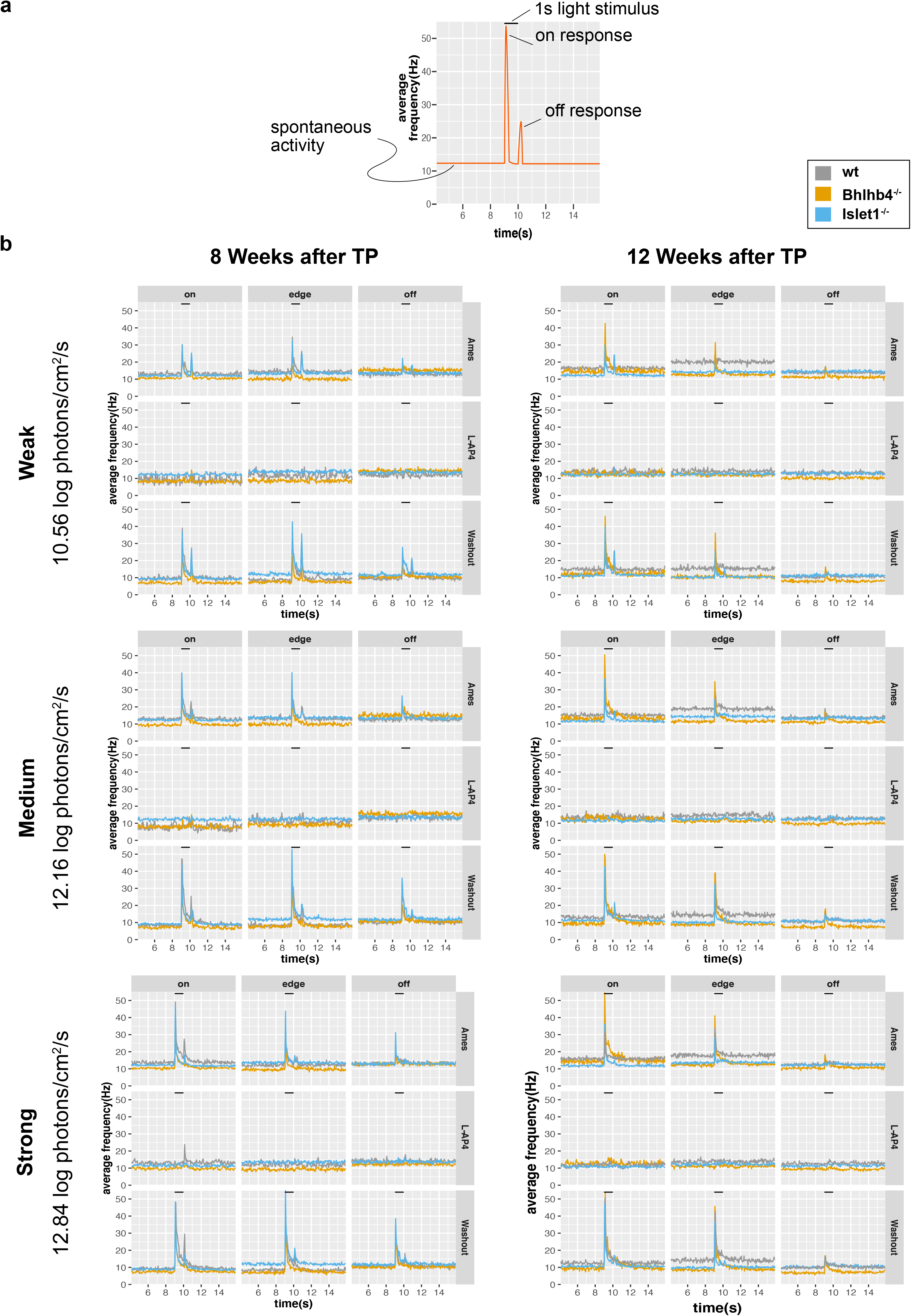
Population averages of recordings with 1 s stimulus. (a) An explanatory illustration. The two peaks at the onset and offset of light stimulation (black bar on top) indicate the increased spiking frequencies as ON and OFF responses and the height of the baseline indicates the spontaneous firing frequency. (b) Population averages of spontaneous activity (the height of the baseline) and light responses (peaks triggered by onset and offset of light stimulus) on different graft areas (on, edge, off) and before, during, and after L-AP4 treatment. The bar on top indicates the 1 s light stimulation. The three rows show recordings at different stimulus strength (weak, medium, strong) and the two columns show responses from samples collected 8 weeks after transplantation (left) and 12 weeks after transplantation (right). Graft genotype is indicated by the color of the trace.

**Supplemental Figure 10.**
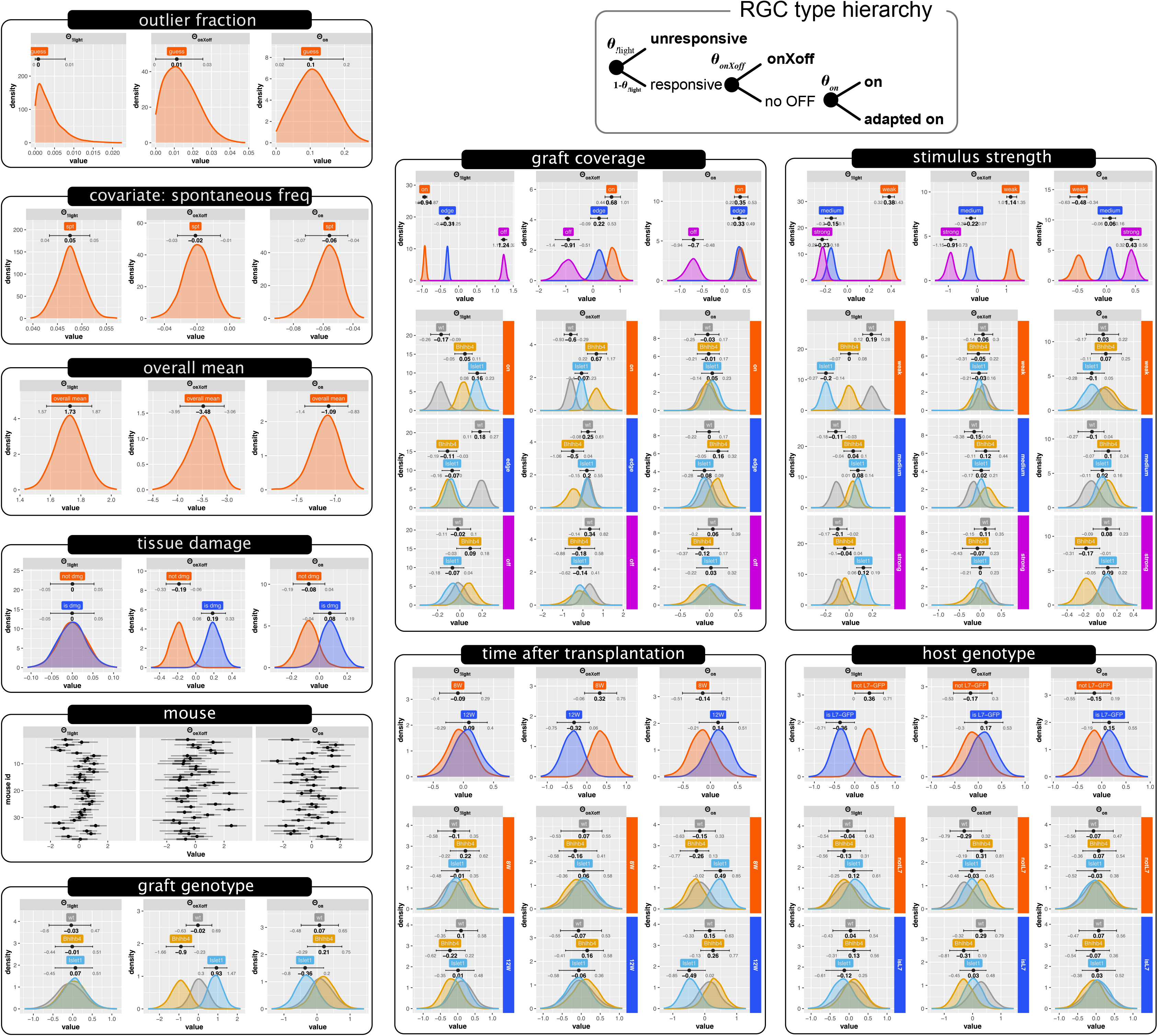
Analysis of RGC response types. The response pattern (ON, OFF, etc.) of individual cells to the 1 s light stimulus through the full before-during-after L-AP4 treatment procedure, and separated RGCs to 4 functional types: being unresponsive to light (!light, the exclamation symbol denotes the logical NOT operator) or being responsive with robust ON response from the beginning (on), with ON response enhanced or emerging later (adapted on), or with response containing the OFF-pathway component (onXoff). These response types were assumed to follow the hierarchy shown in the scheme (RGC type hierarchy) on top. We then used a conditional logit model to estimate the probability of response at branching points (*θ*_!*light*_, *θ_onXoff_, θ_on_*). Graphs show the posterior distribution of parameter with 89% compatibility interval and mode indicated on top. Higher values in the posterior distribution indicate higher values of the probability for that response. Note that because of the model hierarchy, the probability of responsive RGC is estimated as the complement, i.e. probability of no light response, and therefore the probability of light response is higher at lower values of !*light*. Panels on top graphs indicating the response type (!*light, onXoff, on*). Note that although there are four response type (!*light, onXoff, on*, and *adapted on*) there are only three probabilities (*θ*_!*light*_, *θ_onXoff_, θ_on_*) as we are estimating the probabilities at the branching points. Note also that the calculated probabilities do not directly correspond to the ratio of each functional type, as *θ* is calculated at each branching point, excluding the responses from the higher hierarchy. So, the ratio of cells that respond to light is represented by (1 – *θ*_!*light*_) and have a probability *θ_onXoff_* to be of type *onXoff*. The ratio of cells that respond to light and are not *onXoff* type is represented by (1 – *θ*_!*light*_)(1 – *θ_onXoff_*) and have the probability *θ_on_* to be of type *on*. Finally, the ratio of *adapted on* is given by (1 – *θ*_!*light*_)(1 – *θ_onXoff_*)(1 – *θ_on_*). Posterior distributions for interaction terms with genotype are shown below the main effect distributions for graft coverage, stimulus strength, host genotype, and time after transplantation. The horizontal axis, confidence interval, and mode for “guess” represents the fraction [0-1] of data points that are estimated to be generated by a random process, i.e. outliers. The horizontal axes, confidence intervals, and modes for the rest of the parameters are in log odds with higher values indicating higher probability for the respective response category. n=16353 observations, from 5738 cells, collected from 38 mice (9 wt, 13, *Bhlhb4^-/-^*, and 16 *Islet1^-/-^*).

**Supplemental Figure 11.**
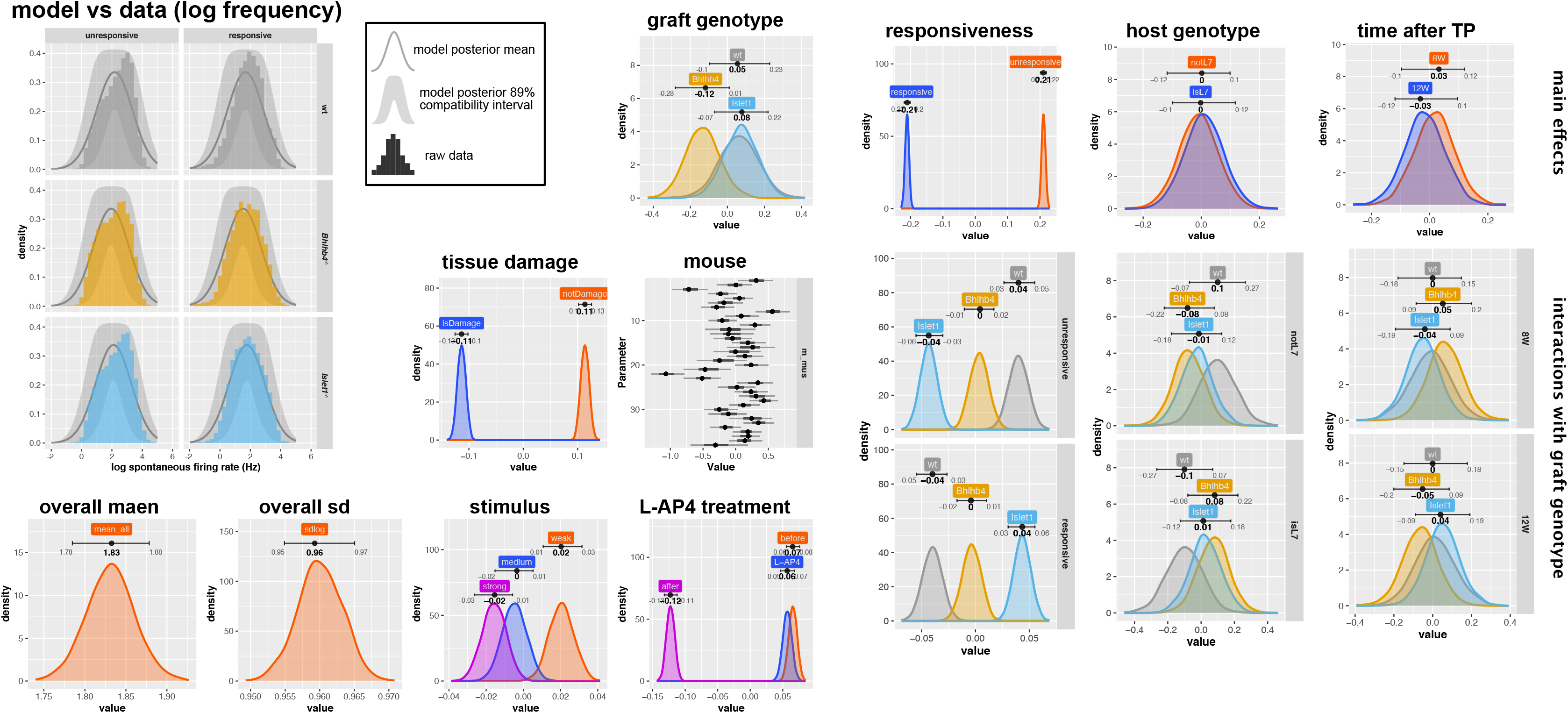
Analysis of RGC spontaneous firing. The spontaneous activity of RGCs was modeled with the lognormal distribution, using the 9 s average spiking rate of cells prior to light stimulation. The log frequency of spontaneous firing follows a normal distribution. We estimated the effect of various parameters on the logmean. Graphs show the posterior distribution of estimated parameters with 89% compatibility interval and mode. Posterior distributions for interaction terms with genotype are shown below the main effect distributions for light responsiveness, host genotype, and time after transplantation. Higher values indicate a higher logmean, and therefore a higher spontaneous activity. n=44213 observations, from 5738 cells, collected from 38 mice (9 wt, 13, *Bhlhb4^-/-^*, and 16 *Islet1^-/-^*).

**Supplemental Figure 12.**
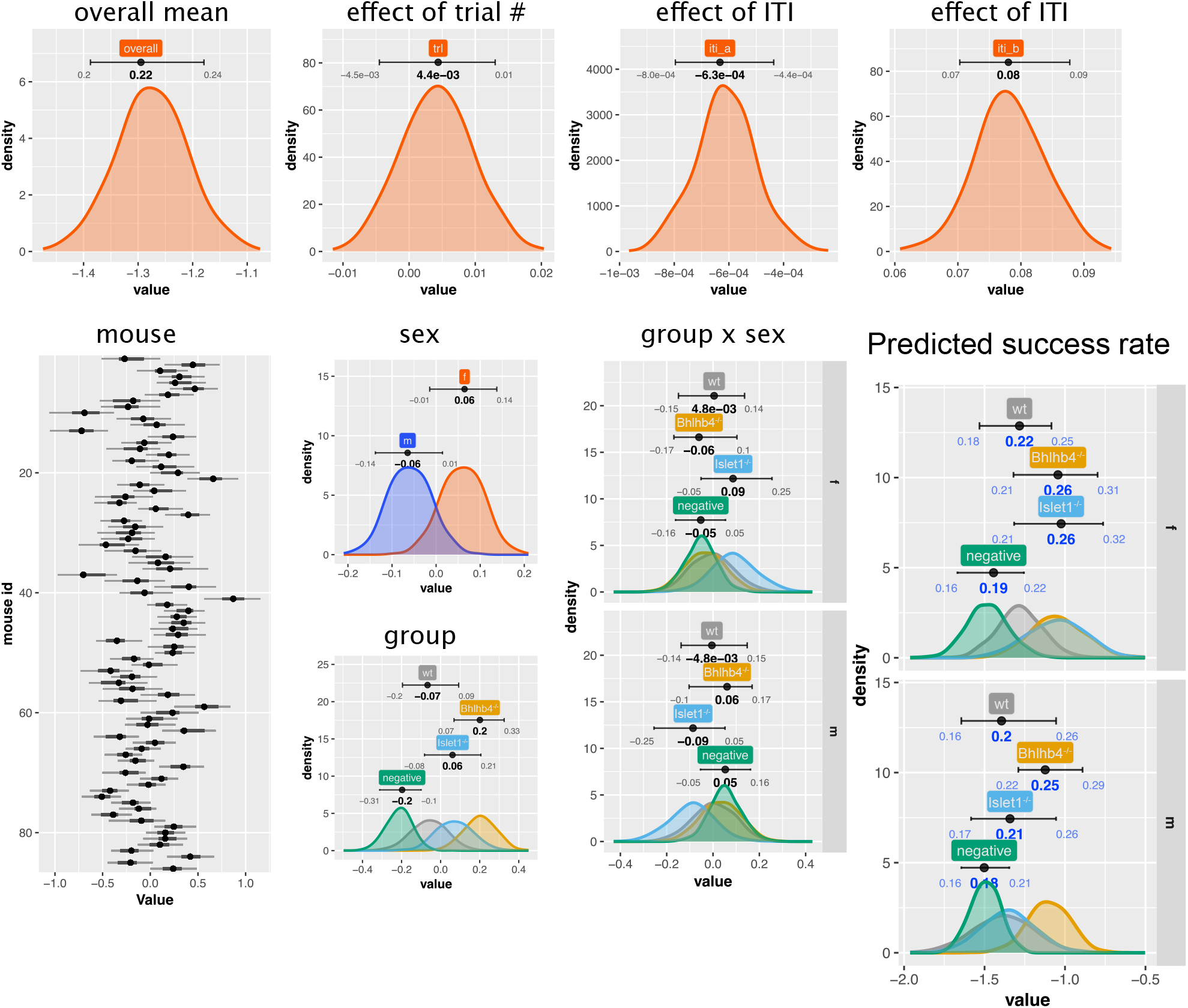
Analysis of light Avoidance Test. The number of success counts *y_i_* was modeled using the binomial distribution (*y_i_* ~ *Binomial*(*θ_i_*, 30)). In addition to group (control, wt, *Bhlhb4^-/-^*, and *Islet1^-/-^*), sex (male or female), interaction of group and sex, and mouse, we also used trial number (*trl*) as a predictor as accumulated experience may improve the performance. Additionally, we included a quadratic term for ITI (*iti*), which is a proxy for the random movements between chambers. Success count increases with ITI as mice that move often can avoid the shock purely by chance. We assume a quadratic relationship as we noticed that the benefit of moving often decreases at high iti as the animal tires. The probability of success is then given by 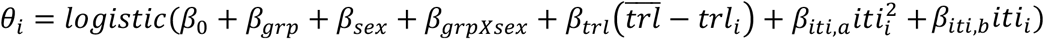, where *β*_0_ is the overall mean. Graphs show the posterior distribution for the model parameters with 89% compatibility interval and mode indicated on top of each posterior distribution. The horizontal axis for “group” (*β_grp_*), “sex”(*β_sex_*), “mouse” (*β_mus_*), “group x sex”(*β_grpXsex_*) represent log odds, as the data was modeled using the logistic function as link function. The “predicted value” indicates the expected value of different grafts transplanted to host of different sex ignoring mouse variation (*β*_0_ + *β_lin_* + *β_sex_* + *β_linXsex_*). The values in the interval and mode for “overall” and “predicted value” are shown converted to the original scale, i.e. *logistic*(*β*_0_) and *logistic*(*β*_0_ + *β_grp_* + *β_sex_* + *β_grpXsex_*) respectively.

## Materials and Methods

### ES/iPS cell lines

Tg(*Nrl*-GFP) is a transgenic iPS cell line expressing GFP under the transcriptional control of *Nrl* promoter^19,33^. ROSA26^+/*Nrl*-CtBP2:tdTomato^ is a *Rosa26* knock-in expressing a CtBP2 and tdTomato fusion protein under the *Nrl* promoter^12^. These ROSA26^+/*Nrl*-CtBP2:tdTomato^ constructs will be referred to as Ribeye-reporter. We also constructed a Ribeye-reporter line on the C57BL/6J-Tg(*Thy1*-GCaMP6f)GP5.5Dkim/J ES cell line (*Thy1*-GCaMP6f;Ribeye-reporter)^34^ Synapse count analysis, and behavior tests were conducted with retinal organoids derived from the Ribeye-reporter ES cell lines. Immunohistochemistry characterization (except for synapse analysis), microarray analyses, and electrophysiology assays were conducted with grafts derived from the Tg(*Nrl*-GFP) lines.

### Generation of Bhlhb4^-/-^ and Islet1^-/-^ cell lines

*Bhlhb4^-/-^* lines were constructed by targeting sites upstream and downstream of the start codon as shown on the schematics in Figure 1a. *Islet1^-/-^* lines were constructed by targeting an upstream site of the start codon in exons 1, and the flanking region at the 3’-end of exon 2. Guide RNAs were designed using the CRISPR design tool (http://crispr.mit.edu). The oligo pairs were cloned into the BbsI-BbsI site in the plasmid containing Cas9 and the single-guide RNA (sgRNA) scaffold (pSpCas9(BB)-2A-Puro, Addgene plasmid IDs: 481398 and 48139), to generate puromycin-Cas9 fusion plasmids co-expressing sgRNAs^35^.

Tg(*Nrl*-GFP);Ribeye-reporter iPS cells and *Thy1*-GCaMP6f;Ribeye-reporter ES cells were transfected with Cas9-sgRNA complexes targeting *Bhlhb4* or *Islet1*, using the mouse ES Cell nucleofector kit (Lonza) and Amaxa Nucleofector II device (Lonza) following manufacturer’s instructions. Transfected cells were treated with antibiotic (0.5-1 ug/mL puromycin) 24 hours after transfection for 48 hours. 4-10 days after antibiotic selection, antibiotic resistant colonies were collected for passage and genotyping. Genotyping was performed by PCR using specific primers to amplify fragments from the wildtype allele and of a shorter fragment from the deleted alleles (Figure 1). The PCR products from genotyping were purified (Macherey-Nagel) and sequenced (Thermo Fisher Scientific) to confirm the deletion. At least 2 or 3 clones were obtained for each KO line.

### Animals

All the experimental protocols were approved by the animal care committee of the RIKEN Center for Biosystems Dynamics Research (BDR) and were conducted in accordance with local guidelines and the ARVO statement on the use of animals in ophthalmic and vision research.

C57BL/6J-Pde6b^*rd1-2/rd1-2J*^/J (JAX stock #004766, hereinafter referred as *rd1*) mice were used as end-stage retinal degeneration model. B6;CBA-Tg(*Thy1*-GCaMP3)6Gfng/J (JAX stock #017893, hereinafter referred as *Thy1*-GCaMP3) mice^36^ were crossed with *rd1* mice to generate *rd1*; *Thy1*-GCaMP3 animals for live imaging using two photon microscopy. B6;FVB-Tg(Pcp2-EGFP)2Yuza/J (JAX stock #004690)^37^ crossed with *rd1*-2J (hereinafter referred as *rd1*;*L7*-GFP) were used for host graft synapse analysis and electrophysiology.

### Transplantation of 3D retinas

Maintenance, differentiation and optic vesicle structure preparation for transplantation of ES/iPS cell derived retinas were as previously described^8^. Briefly, optic vesicle structures (DD10-15) were cut to small pieces (around 400 μm × 500-800 μm), on the day of transplantation, and inserted sub-retinally into the eye, at approximately 0.5-1 mm from the disk at the ventro-temporal side, of the 8-12 week old *rd1* and *rd1;L7*-GFP mice with 1 mM VPA using a glass micropipette with a tip diameter of approximately 500 μm. Indomethacin (10 mg/L) was added to the drinking water of all transplanted mice starting on the day of transplantation.

### RT-PCR of retinal organoids

Total RNA was extracted from the differentiated retinal organoids using TRIzol reagent (Invitrogen) and cDNA was synthesized using Oligo(dT) primers and SuperScript III reverse transcriptase (Invitrogen). Target genes were amplified using SapphireAmp Fast PCR Master Mix (TaKaRa) with the specific primers detailed in the supplemental materials.

### Microarray analysis

Total RNA was extracted from the differentiated retinal organoids (5 retinal organoids for each condition) using TRIzol reagent (Invitrogen) and was purified using RNeasy kit (QUIAGEN) following manufactures’ instructions. RNA quality check, concentration, and data collection using the SurePrint G3 Mouse GE microarray 8×60K Ver. 2.0 (Agilent technologies) were performed by Hokkaido System Corporation (Hokkaido, Japan), according to Agilent protocols. The array contained 56,605 probe sets covering 37,074 genes. Data processing, analyses, and visualization was done in R^38^ using custom scripts. Probes below background level were discarded. For Gene ontology (GO) analysis, GO annotation for the microarray (downloaded from Agilent’s site) were correlated with GO annotations in the Mouse (GO.db: a set of annotation maps describing the entire Gene Ontology. R package version 3.6.0.).

### Immunohistochemistry

After mice were sacrificed by cervical dislocation, the eyes were enucleated and immediately fixed with 4% paraformaldehyde (PFA) for an hour at 4 °C. The eyes were then washed with phosphate buffered saline (PBS), followed by the removal of the cornea, iris, and lens under a microscope. For cryosection preparation, the eyes were infiltrated with 30% sucrose overnight at 4°C and embedded in optimum cutting temperature (OCT) compound (4583, Sakura Finetek Japan, Tokyo) and stored at −30 °C. Coronal plane cryo-sections of 10-14 μm thickness were made with a cryostat (Leica CM3050S). After two rounds of PBS wash, sections were exposed to 0.1% Triton X-100 in PBS (vol/vol) for 30 min to permeabilize cellular membranes. Cells were then blocked with blocking solution for 1 h at room temperature. Primary antibodies were diluted in blocking solution and incubated with samples overnight. Samples were then rinsed three times with 0.05% Tween-20 in PBS (vol/vol) and incubated with the appropriate secondary antibodies for 1 h at room temperature and counterstained with DAPI (Life Technologies).

For free-floating immunostaining, 50 μm thick retinal flat mount sections were collected into 12-well tissue culture plates containing cold PBS (5-8 sections/well) and blocked with staining solution (3% Triton X-100 and 1% BSA-PBS) at 4°C overnight. The samples were then incubated with primary and secondary antibodies in staining solution at 4°C for three to four days and two days, respectively, and mounted with 2,20-thiodiethanol (Sigma) or VECTASHIELD (Vector). All staining procedures were performed with gentle mixing on a rocker.

Images were acquired with a Leica TCS SP8 confocal microscope and reconstructed in 3D Imaris Microscopy Image Analysis Software (http://www.bitplane.com/) and Fiji (ImageJ).

### Quantification of graft rods

The number of rods in grafts was quantified from Z-stack images of flat mounted retinas from mice sampled roughly 60 days (8-9 weeks) post transplantation. Orthogonal sections at coordinates that intersected grafts were resliced in Fiji and the graft area was traced manually.

Using grafts with the *Nrl*-GFP reporter, rod photoreceptors can easily be identified by the GFP signal, as well as by their size, localization and DAPI signal pattern. We delineated the area of the graft manually and counted the number of cells in the graft (DAPI^+^ cells) and the number of rod photoreceptor cells (GFP^+^ cells) using a Fiji macro. Figures 2a-d show a typical image of this quantification process. Fiji macro scripts are provided in github repository (https://github.com/matsutakehoyo/KO-graft).

### Quantification of graft inner cells

The numbers of inner cells, Calbindin^+^, Calretinin^+^, PKCα^+^, and Chx10^+^ cells in the grafts were counted manually from retinal sections of mice sampled roughly 30 days (4 weeks) post transplantation. The total population of graft inner cells was determined by elimination. First, host area, characterized by the laminar structure of the host inner nuclear layer, was identified. Secondly, graft photoreceptors were identified by *Nrl*-GFP, nucleus morphology, and local distribution within rosettes. The remaining cells were considered inner cells and the total number of inner cells and the number of cells positive for a particular marker were counted.

### Host-Graft Synapse Count

For the host-graft synapse count, *Thy1*-GCaMP6f;Ribeye-reporter ESC-retina was transplanted to *rd1*;*L7*-GFP mice and the eyes were collected at DD50 (approximately 5 weeks after transplantation). Regions containing part of the graft rosettes were cropped and the number of *L7*-GFP positive bipolar cells were manually counted by one observer in a blind manner. A synapse was defined as a pairing of the tdTomato signal from the RIBEYE reporter at the graft rod presynaptic terminal (*Thy1*-GCaMP6f;Ribeye-reporter) and the *Cacna1s* signal (stained) at the post synaptic terminal. Importantly, only *Cacna1s* signals at the dendritic tips of *L7*-GFP cells were counted, thus ensuring that counted synapses were strictly between graft photoreceptors and host bipolar cells. Furthermore, the pairing of synaptic markers was judged from three orthogonal sectional views (X-Y, X-Z, and X-Z) independently to avoid any optical artifacts.

### Electrophysiology: multi-electrode array (MEA) recording

Micro electroretinogram (mERG) and RGC spiking activity were recorded with the MED64 system (Alphamed Scientific Inc., Osaka, Japan) with a standard 8×8 probe (MED-P515A) as previously described^13,14,39,40^. Following 1-3 days of dark adaptation, mice were anesthetized with sevoflurane (Abbott Japan, Osaka, Japan) and sacrificed by cervical dislocation. After enucleation, the transplanted retina was carefully isolated and vitreous was gently but thoroughly cleaned. The position of the graft was visually targeted in between the optic nerve disc and the transplantation entry site to be mounted over the probe with the ganglion cell side against the electrodes. A dim LED light source with the peak wavelength at 690 nm was used during sample preparation. The flat-mounted retinas were constantly supplied with warmed (34 ± 0.5°C), carbonated (95% O_2_ and 5% CO_2_) Ames’ medium (A1420, Sigma-Aldrich) perfused at 3-3.5 mL/min. Opsinamide (10 μM; AA92593, Sigma-Aldrich) was added in the perfusion medium to suppress the melanopsin-driven RGC light responses during recording. Retinas were kept in the dark, allowed to recover in the chamber for at least 20 min before the first set of stimulation while spontaneous spikes were monitored from the beginning. Full-field stimuli with intensities of 10.56 (weak), 12.16 (medium) and 12.84 (strong) log photons/cm^2^/s (equivalent to approximately 0.03 to 6.5 lux or 1.7×10^2^ to 3.4×10^4^Rh*/rod/s) at the focal plane of probe were generated with a white LED (NSPW500C, Nichia Corp., Tokushima, Japan) without background illumination. An electronic pulse generator (DSP-420, Dia-Medical System, Tokyo, Japan) was used to control the LED intensity under the external command from MED64 system using scheduled TTL pulses to turn on and off the LED. Pulse stimuli of 10 ms were used to elicit the mERGs and 1 s stimuli were used to separate the RGC response types. In our standard protocol, three-time serial recordings were repeated with an interval of 1 min, and a single recording time was 10 s for the 10 ms stimuli and 20 s for the 1 s stimuli. Data were collected at 20 kHz sampling rate and filtered at 1 Hz (10 ms stimuli) or 100 Hz (1 s stimuli) online. Recordings were repeated before and after 9-*cis*-retinal (100 μM for 10 min incubation in the recording probe; R5754, Sigma-Aldrich) replenishment, as well as in the presence and absence of L-AP4 (20 μM; 016-22083, Wako) blockade to confirm the reproducibility.

Graft locations were manually mapped on a 26×26 grid overlaid on the MEA electrode array. Our MEA probe consisted of 50×50 μm^2^ channels separated by 100 μm in an 8×8 array. The size of a channel corresponds to a square in our 26×26 map, and the separation between channels correspond to two squares. On each square we mapped the presence or absence of graft (on, off) and applied a morphology transformation to dilate the graft area to define graft edges. In addition to graft location, areas where tissue damage was apparent or covered by the optic disc were marked as “damage” assuming these areas would have some kind of impairment to respond.

mERG traces were processed and analyzed in R^38^. A band-pass Butterworth filter (1 to 50 Hz) was applied to traces to remove low frequency fluctuations and high frequency jitter. Local minima within 55 ms from light stimulation were flagged as a-wave, and local maxima within 150 ms from light stimulation were flagged as b-wave. The a-wave amplitude was calculated from the baseline, and the b-wave amplitude was calculated from the a-wave, or from the baseline when the a-wave was not detected. Replicates from three repeated stimulations were averaged.

RGC spike sorting was conducted using the CED Spike 2 software (version 7.2). Recordings taken at the same stimulus intensity were processed together by merging recordings taken before 9-*cis*-retinal treatment and before, during and after L-AP4 treatment, to follow spike trains from the same set of cells across the different conditions. Recording traces were first cleaned up by applying a Butterworth band-pass filter (200 to 2800 Hz) and DC offset removal. The spike templates were generated for each channel independently, based on the automatic offline template formation and spike matching algorithm of Spike 2 with a few minor modifications. Spikes with amplitudes smaller than 64 μV (negative) were excluded; this threshold level was about 10-sd of recording without retina and 4-sd of recording with robust spontaneous spiking of degenerated retina. As the whole recording procedure lasted for more than 3-4 hours, sometimes the spike amplitudes were found amplified over time. A 5% tolerance for maximal spike amplitude change was therefore included in the template matching process so that each spike would be slightly resized (±5%) to see if its shape matches any established template. This process allowed us to isolate the individual spike sources (cells) with characteristic spike size and shape from each sample for the given stimulus intensity. The average number of cells detected per channel was 2.4 ± 2.9 (mean±sd) for 1 s stimulus recordings (4 ± 2.5 when excluding channels with no cells), and 2.4 ± 2.8 for 10 ms stimulus recordings (4 ± 0.26 when excluding channels with no cells).

The assignment of light responsiveness (10 ms stimuli) or light response types (1 s stimuli) was determined arbitrarily by comparing the spiking patterns following light stimulus to the basal frequency. Cells with basal spiking frequency lower than 1 Hz were considered inactive and excluded from the following analysis. In contrast, active cells with basal spiking frequency higher than 3 Hz were determined as “spontaneous firing”. In the 10 s recordings cells were simply defined as light responsive if showing 2-fold or more spiking than resting state within 500 ms after the 10 ms stimuli. The response types to 1 s stimulus for each cell in each condition were first described as non-responsive, transient ON, sustained ON, delayed ON, ON suppression, OFF, ON-OFF and hypersensitive separately (see also Iraha et al., 2018), by comparing the onset and termination timing of their spiking frequency changes to the stimulation. Cells were then grouped as “!light” (unresponsive), “on”, “adapted on” or “onXoff’ types based on their response patterns across all three conditions (before, during and after L-AP4). In principle, “on” cells showed L-AP4 sensitive ON (including sustained ON) responses throughout the whole recording, while “adapted on” cells had their ON responses (mostly sustained ON) seen only after recovery from L-AP4. Although cells with OFF responses were also included, these rarely found “onXoff” type mostly consists of cells showing ON and OFF responses that were both sensitive to L-AP4 blockade, suggesting an ON-dependent OFF pathway involvement in the synaptic inputs to these cells. Note that delayed ON and hypersensitive responses were seldom observed and proved independent to the synaptic inputs, therefore assigned to the “!light” unresponsive type. For both recordings with 10 ms and 1 s stimuli, responses that were not consistent across the three replicate recordings were disregarded. For the spontaneous firing analyses, we averaged the spiking activity prior to light stimulation (9 s duration from the 20 s recording) of the three replicate recordings.

### Electrophysiology: two photon imaging

*rd1*;Thy1-GCaMP3 mice transplanted with retinal sheets (wt, *Bhlhb4^-/-^, Islet1^-/-^*) were sacrificed at various time points after transplantation (7-26 weeks) by cervical dislocation, after at least 12 hours of dark adaptation prior to recordings. Areas away from the graft (designated as “off”) served as negative controls, as the grafted cells only covered a small fraction of the retina. Age matched *Thy1*-GCaMP3 animals were used as positive controls (“normal”). Eyes were enucleated, and the retina were flat mounted using a mesh covered anchor with the ganglion cell layer on top. Retinas were constantly supplied with warmed (28 ± 0.5°C), carbonated (95% O_2_ and 5% CO_2_) Ames’ medium (A1420, Sigma-Aldrich) perfused at about 10 mL/min. Sulforhodamine 101 (500 nM; S7635 Sigma) was added in the perfusion medium to confirm ganglion cell viability and to help locate graft. The ganglion cell layer plane was identified using sulforhodamine counterstaining and the basal fluorescence signal from GCaMP3. Images were acquired with an upright multiphoton laser scanning microscope (Olympus, FV1000MPE) using a water-immersion 25x objective lens (Olympus, XLPLN25XWMP). The light source was a Mai Tai femtosecond-pulse laser (Spectra-Physics) running at 920 nm. The graft position was identified by the fluorescence of the graft (*Nrl*-GFP). Images were acquired at 0.066 s/frames on an area of 254.46 μm x 254.46 μm at 128 px x 128 px resolution for approximately 152 s (2300 frames). Retinas were recorded for at least a minute before a brief pulse light stimulation, at around 90 s, to allow transient fluorescent changes to equilibrate. The light source for the stimulus was a homogenous green light (λ=505 nm LED passed through a collimator lens) of 5.78×10^-13^ W/μm^2^ which would be equivalent to 1.28×10^6^ Rh*/Rod/s in a normal mouse retina. Images were processed in Fiji (ImageJ) using custom scripts by Z-projecting time series images and extracting ROIs (Regions Of Interest) using the analyze particles option. The fluorescence intensity of each ROI along the time axis was then exported to be processed in R. We defined a Light Response Index (LRI) as below, to compare fluorescence intensity changes before and after light stimulation.

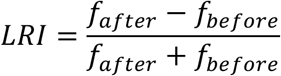

*f_before_* and *f_after_* represent the fluorescence changes before and after light stimulation, which are calculated relative to the baseline fluorescence.

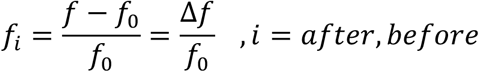

Finally, the light response of each ROI was weighted by the total area of detected ROIs and aggregated to obtain the light response for each recording.

### Behavior: Light Avoidance

The shuttle-avoidance system (CompACT SAS/W, Muromachi Kikai) was used to evaluate the light responsiveness of transplanted mice as previously described^12^. Briefly, a dark-adapted mouse is place in a sound- and light-insulated box with two chambers able to freely move between chambers. The mouse is then presented with stimulus at random intervals at one of the chambers. If the mouse is stationary the stimulus is proceeded by a mild electric shock (0.08mA) delivered on the floor bars. The mouse can avoid the shock if it moves away from the chamber within 5 s of stimulus presentation. Each experiment consisted of 30 such trials, and the number of successful avoidances is recorded. Mice able to sense light (3 cd) can learn to use light as a warning and avoid the shock. Mice may also increase their success rate by frequently moving between chambers, so the number of times the mouse moves before light stimuli (ITI, inter trial interval) is used as a proxy to account for this behavior. Initially mice were trained with a dual sound and light stimuli, and only mice that successfully learn to respond to the dual stimuli were subsequently tested with a light stimulus only. Mice were tested daily, except for weekends, until their response appeared to plateau.

### Statistical analysis

We used full Bayesian statistical inference with MCMC sampling for statistical modeling using Rstan (Stan Development Team. 2017. *RStan: the R interface to Stan*. R package version 2.16.2. http://mc-stan.org). We estimated population effects, such as the effect of different graft genotype (wt, *Bhlhb4^-/-^, Islet1^-/-^*), group effects such as the effect of different treatments or conditions, inter-mouse variation, and others, using linear regression models. In this framework, an observation from a particular set of treatments or conditions, coming from a certain mouse in a certain group is simultaneously used to estimate the effect of that group, mouse, and population. Our estimates, therefore, do not represent a simple pooling of data for a particular set of predictor combinations. Instead, linear regression analyses can isolate the effect of predictors while considering the data as a whole.

During the course of the study it became apparent that some of the outcomes could be potentially be influenced by the sex of the host animal, and so whenever possible we included the host sex as a predicter, this however was not feasible in all of the analyses because of logistics and in such cases the analysis was conducted on males only discarding the data for the females.

Along with the estimation of main effects, the overall effect of predictors, we estimated interactions, the conditional effects that manifest under a particular combination of predictors in addition to main effects. We mainly estimated interactions of genotype with other predictors (e.g. interaction of genotype and sex, denoted genotype x sex), in order to characterize the genotype effect in more detail. Whenever an interaction term was considered, we present the main effects and interaction effects together, as interaction terms must be considered alongside their main effects.

Posterior distribution of parameters of interest, which show the probability for the value of the parameter given the data, are shown with 89% compatibility intervals (confidence intervals) and mean indicated on top. This interval was chosen as it gives more stable estimates than the traditional 95% interval and highlights the arbitrariness of interval limits. These marginal distributions of parameters do not necessarily indicate the relationship between parameter values as parameters maybe correlated. In order to judge if there is a difference between parameters, we must look at the joint distribution of parameters, i.e., the distribution of the difference between parameter values. Whenever a difference is indicated we therefore indicate the fraction of the area over zero, which indicates the confidence that we have that there is a difference.

Many of the models use a link function (logistic, exp, etc.) and so the interpretation of the results is not intuitive. In order to facilitate interpretation, distribution of the posterior predictions is show in the supplemental data with compatibility intervals translated to the original data scale. Stan scripts for the models are provided in a github repository (https://github.com/matsutakehoyo/KO-graft).

### Analysis of Nrl-GFP fluorescence of retinal organoids

Image analysis was conducted using EBImage^41^ and statistical modeling was implemented using Stan^42^ in R^38^. Acquired fluorescence images were recorded as RGB images. Pixels containing the GFP signal were identified from background by thresholding using Otsu’s method, and the mean signal and background signal of the green channel was calculated for each image. Signal and background were also calculated for the blue channel to account for differing recording conditions by normalizing the fluorescence intensity on the blue channel. The relative fluorescence intensity *y* is thus expressed as

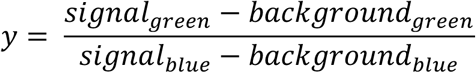

The relative fluorescence intensity was fitted with a modified Gompertz growth curve^43^ with the Gamma distribution as likelihood, as relative fluorescence is continuous value that is always positive. The modified Gompertz growth curve, is defined by three parameters representing maximum fluorescence intensity *A_line_*, maximum rate of fluorescence increase *μ_line_*, and onset of fluorescence increase *λ_line_* (see Supplemental Figure 1). Thus the fluorescence intensity for a *line* (wt, *Bhlhb4^-/-^* or *Islet1^-/-^*) on time *day* is

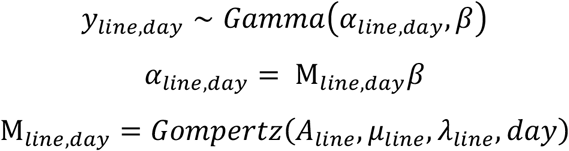

Where *α* and *β* represent the shape and rate parameters of the Gamma distribution, and *M* represents the average fluorescence intensity.

### Graft cell number

The number of photoreceptor cells in the graft and the number of cells positive for PKCα, Chx10, Calbindin, and Calretinin, within the inner cells of the graft were modeled with the Binomial distribution, as these represent discrete count data with a binary outcome. Thus, the number of positive cells *y_i_* for a particular marker in a given sample *i* is given by

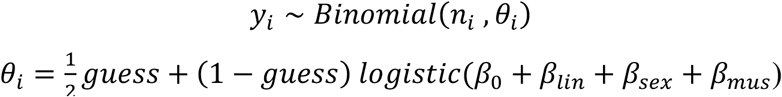

Where *n_i_* represents the total number of cells present in the sample and *θ_i_* ∈ [0,1] represents the probability of cells being positive for the marker. We implemented robustness in our model using a *guess* ∈ [0,1] parameter. In this model *θ_i_* is given by two parts, a purely random part (*guess*)and the logistic regression of the predictors (*logistic*(*β*_0_ + *β_lin_* + *β_sex_* + *β_mus_*)). A large *guess* value indicates the data contain many points not conforming to the regression. For the logistic regression we considered three predictors: the genotype of the cell (either wt, *Bhlhb4^-/-^*, or *Islet1^-/-^*) *β_gen_*, the sex of the host *β_sex_*, and the mouse *β_mus_* from which the sample was derived. *β_sex_* was omitted from the analysis of photoreceptor cell number as samples consisted of only males. *β*_0_ represents the overall mean and the effect of a particular predictor is modeled as a deviation from the overall mean with a sum-to-zero constraints for each predictor (∑ *β*_[*predictor*]_ = 0). Generic weakly informative priors were used for all parameters.

### Number of synapses

Bipolar cells typically make multiple synapses to photoreceptor cells. We therefore counted the number of synapses per bipolar cell. The collected data contained a large zero count peak, indicating that there was a large population of bipolar cells that do not make connections at all, and a second population that makes multiple synapses as shown in Figure 3j. The data was therefore analyzed with a zero-inflated Poisson model, assuming that the number of synapses draws both from a Bernoulli distribution with probability *θ* and a Poisson distribution with mean *λ*. In this case whether a synapse is formed at all is modelled as a Bernoulli process (either success or failure) and the number of synapses per L7-GFP bipolar cell is modelled with the Poisson distribution. A host bipolar cell (L7-GFP) therefore has a probability *θ* to form a chemical synapse with graft photoreceptor cells, and the number of connections it makes is described by *λ*. The probability function of synapse numbers *p*(*y_n_|θ_n_, λ_n_*) is given by

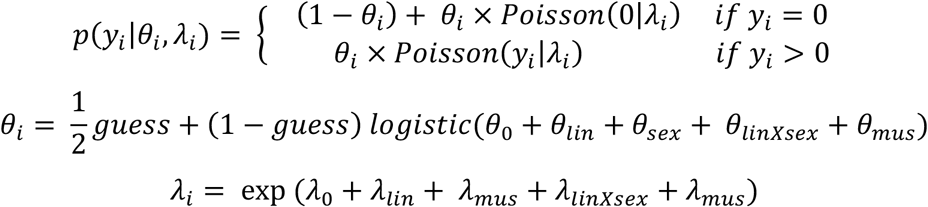

Where *θ_i_* represent the probability of non-zero (probability of synapse) of the *i*-th data, and *λ_i_* represent the average number of synapses, with the *guess* parameter similarly implemented as in the cell type analysis. Predictors include *lin* (wt, *Bhlhb4^-/-^*, or *Islet1^-/-^*), *sex*, and *mus* (mouse), as well as the interaction between *lin* and *sex* (*linXsex*). Generic weakly informative priors were used for all parameters (*Normal*(0,1)).

### Two photon imaging

The light response index was analyzed with a Gamma hurdle model, a mixture of response probability and response amplitude. Whether a response is detected or not (>0) is modelled as a Bernoulli process and the amplitude of the response is modelled with the Gamma distribution. The probability function of response amplitude *p*(*y_n_|θ_n_, λ_n_*) is given by

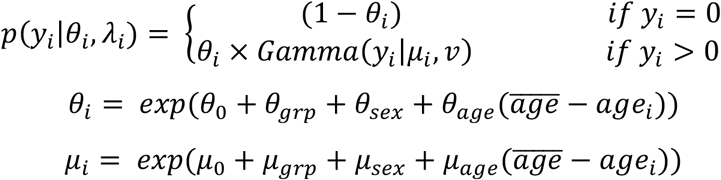

Where *θ_i_* represent the probability of non-zero (probability of response), and *μ_i_* represent the average response amplitude. Predictors include *grp* (wt, *Bhlhb4^-/-^, Islet1^-/-^*, off, normal), *sex*. Fruthermore, and *θ_age_* and *μ_age_* account for the varied sampled times. Generic weakly informative priors were used for all parameters.

### b-wave analysis (10 ms stimulus)

Field potential recordings from individual channels in the MEA probe were processed individually. First, positive peaks within 120 ms of the light pulse were selected as candidate b-wave peaks using a custom program in R. We then calculate the standard deviation of the baseline (2 s prior to the light stimulation) for each trace and channels with a peak amplitude higher than three times the baseline standard deviation were considered to have detected a b-wave. We obtained 12736 recordings from 38 animals transplanted with retinal organoids (excluding the channels assigned to be covered by “damage” retinal areas). We then modeled the probability of observing a b-wave in each channel under different conditions. The probability of observing a b-wave is then given by

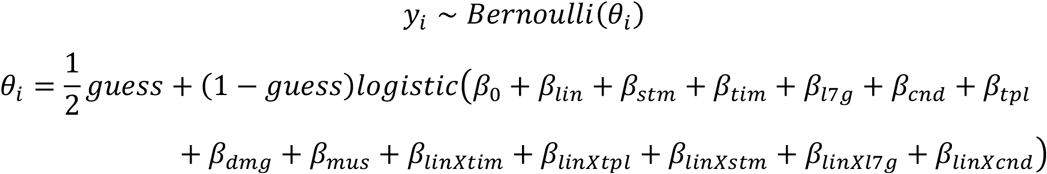

Where *y_i_* is either 0 (no b-wave) or 1 (b-wave) and *θ_i_* is the probability of observing a b-wave in that channel. *β*_0_ represents the overall mean and the effect of different predictors were calculated as a deviation from this mean with the sum-to-zero constraint on each of the predictors (∑ *β*_[*predictor*]_ = 0). Table 1 shows the meaning and value of each of the predictors. Generic weakly informative priors were used for all parameters (*Normal*(0,1)).

**Table 1.** Predictors for analysis of b-wave.

| Predictor | meaning | Value |
| --- | --- | --- |
| $\beta_{lin}$ | Line genotype | wt, *Bhlhb4*<sup>-/-</sup>, *Islet1*<sup>-/-</sup> |
| $\beta_{stm}$ | Stimulus strength | weak, strong |
| $\beta_{tim}$ | Time after transplantation | 8 weeks, 12 weeks |
| $\beta_{l7g}$ | Host genotype | *rd1*, *rd1*;L7-GFP |
| $\beta_{cnd}$ | Condition | before L-AP4, L-AP4, after L-AP4 |
| $\beta_{tpl}$ | Graft topology | on graft, edge of graft, off graft |
| $\beta_{dmg}$ | Tissue damage | no damage, damage |
| $\beta_{mus}$ | Mouse | Mouse id 1...38 |
| $\beta_{lin \times tim}$ | Interaction of line and time | (wt, *Bhlhb4*<sup>-/-</sup>, *Islet1*<sup>-/-</sup>) x<br>(8 weeks, 12 weeks) |
| $\beta_{lin \times tpl}$ | Interaction of line and topology | (wt, *Bhlhb4*<sup>-/-</sup>, *Islet1*<sup>-/-</sup>) x (on, edge, off) |
| $\beta_{lin \times stm}$ | Interaction of line and stimulus | (wt, *Bhlhb4*<sup>-/-</sup>, *Islet1*<sup>-/-</sup>) x (weak, strong) |
| $\beta_{lin \times l7g}$ | Interaction of line and host | (wt, *Bhlhb4*<sup>-/-</sup>, *Islet1*<sup>-/-</sup>) x (not L7, L7) |
| $\beta_{lin \times cnd}$ | Interaction of line and condition | (wt, *Bhlhb4*<sup>-/-</sup>, *Islet1*<sup>-/-</sup>) x<br>(before, L-AP4, after) |

### RGC response (10 ms stimuli)

The spikes from RGC upon the 10 ms stimuli allow us to examine the correlation of b-wave and RGC response. Following spike sorting and response assignment, we modeled the probability of each spike train to be light responsive, i.e. the probability of each RGC to respond to light.

We obtained 30167 observations, from 6827 cells, collected from 38 animals transplanted with retinal organoids. The number of observations roughly corresponds to the number of cells multiplied by six, for the two light stimulus condition (weak and strong) and the three L-AP4 treatment conditions (before, L-AP4, after). As the outcome is dichotomous (either light responsive or not) we used a similar Bernoulli model as in the b-wave analysis with some modifications. In addition to the predictors in the b-wave model, we used the spontaneous frequency rate of each cell, and the b-wave amplitude recorded on that channel as covariates. The probability of an RGC to respond to light is given by

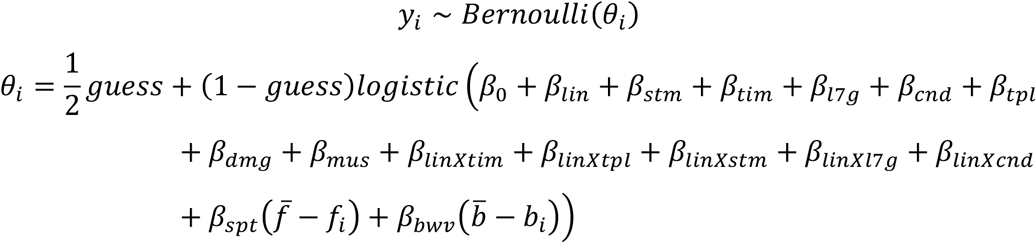

Where *y_i_* is either 0 (unresponsive) or 1 (light responsive) for a particular cell *i* and *θ_i_* is the probability of observing a light response in that cell. *β_spt_* is a coefficient for the effect of the frequency of the spontaneous activity, 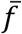 is the mean spike frequency of all the data, and *f_i_* is the spontaneous spiking frequency of the cell *i*. Similarly, *β_bwv_* is a coefficient for the effect of the b-wave amplitude on that channel, 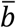 is the average b-wave amplitude, and *b_i_*. Generic weakly informative priors were used for all parameters (*Normal*(0,1)).

### RGC response (1 s stimuli)

The 1 s stimulus allow us to analyze the light response in more detail taking into account different response subtypes and the response patterns under different conditions. For these analyses we used responses from 16353 observations, from 5738 cells, collected across 38 animals transplanted with retinal organoids. The number of observations is roughly three times of the cell numbers for there are three different stimulation conditions.

We categorized response patterns and modeled recorded data with a conditional logit model, assuming that response categories are hierarchical. For the light response we assumed that cells are either responsive or not responsive to light (!light). Of the responsive cells, i.e. excluding non-responsive cells, we assumed that the response either intersects the off pathway (“onXoff’ type) or not. Finally, responses that are independent of the OFF pathway are divided into “on” type or “adapted on” depending on whether they show some sort of potentiation (light response appearing only after several flashes or L-AP4 treatment) or not. Similarly, the spontaneous firing frequency before light stimulation was divided into four categories: <6Hz, 6-12Hz, 12-18Hz and >18Hz to analyze the spontaneous firing. A diagram of the category hierarchy is provided in Supplemental Figure 10. The light response *y_i_* ∈ {! *light, onXoff, on, adp_on_*} can therefore be described as follows

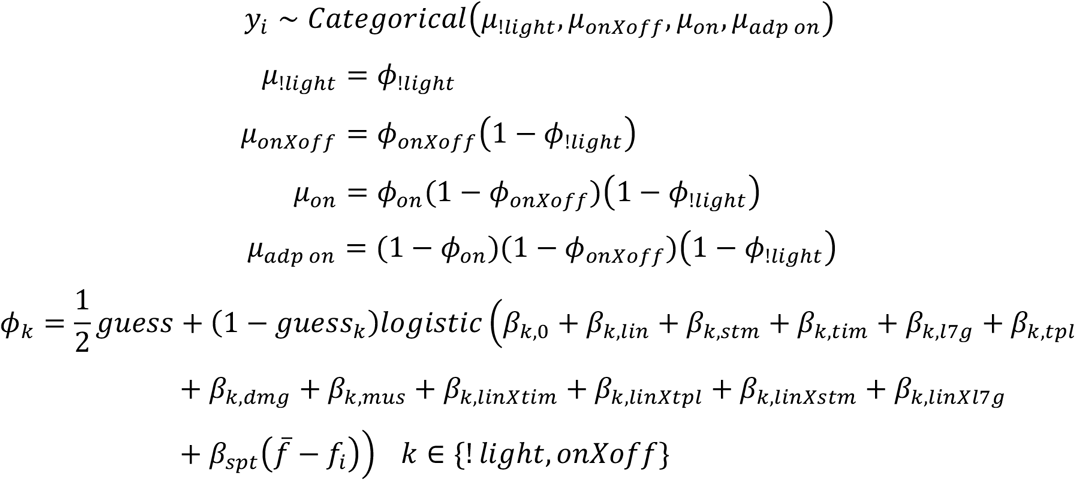

Where *μ* represents the probability of each response category, adding up to 1 (∑*μ* = 1). Note that *ϕ_k_* correspond to the response probability excluding previous categories and therefore does not add up to 1. So *ϕ*_!*light*_ represents the probability of cells not to respond to light, *ϕ_onXoff_* represents the probability of light responding cells to be *onXoff*, and *ϕ_on_* correspond to the probability of non *onXoff* light responsive cells to be *on. β*_*k*,0_ represents the overall mean probability of response category *k*, and the effect of different predictors were calculated as a deviation from this mean with the sum-to-zero constraint on each of the predictors (∑ *β*_[*k,predictor*]_ = 0, *k* ∈ {! *light, onXoff, on*}). Table 2 shows the meaning and value of each of the predictors. Generic weakly informative priors were used for all parameters (*Normal*(0,1)).

**Table 2.**
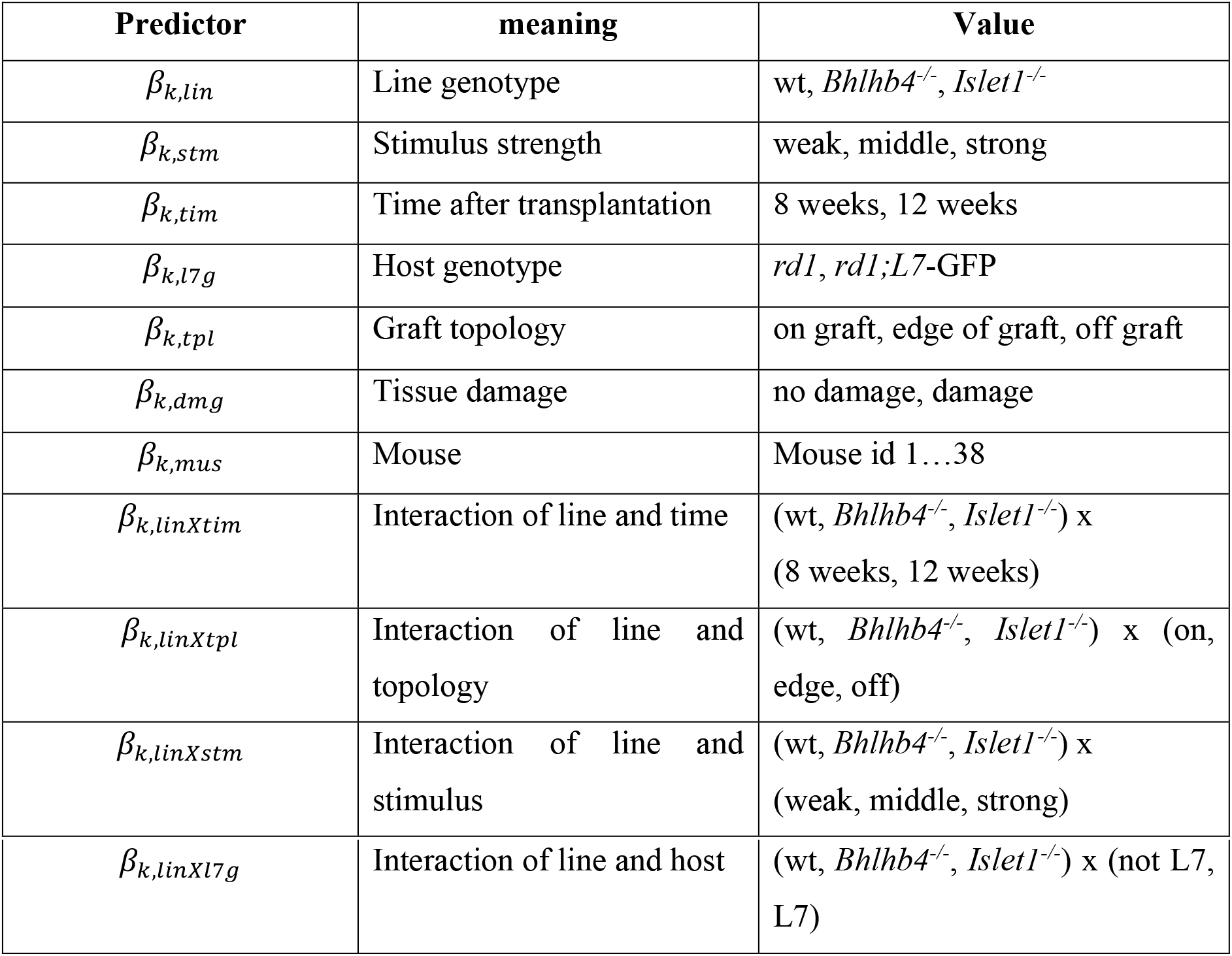
Predictors for conditional categorical model.

### RGC spontaneous firing (9 sec recording before 1sec stimuli)

We noticed that there was an inverse correlation between light responsiveness and the spontaneous firing frequency. We therefore analyzed the distribution of spontaneous firing, by calculating the spontaneous firing rate before light stimulation (9 sec). The distribution of spontaneous firing rate closely follows a lognormal distribution, as shown in figure 6 and supplemental figure 11. We therefore modeled the influence of parameters on the mean log spontaneous firing. For the spontaneous firing analysis, we used 44213 observations from 5738 cells collected across 38 animals transplanted with retinal organoids. The number of observations in this analysis is higher than the number in the light response analysis, as we consider the spontaneous activity before, during, and after L-AP4 treatment, in addition to the different stimuli.

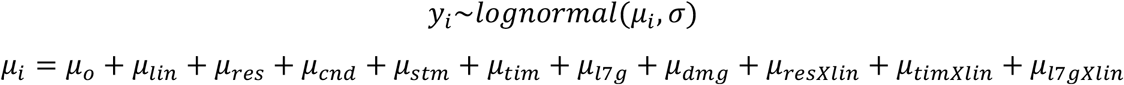

where *y_i_* is the log spontaneous firing ratefor a particular observation *i.μ_o_* and *σ* are the overall logmean and logsd of the lognormal distribution. We assume a common *σ* to simplify the model. The effect of different predictors were calculated as a deviation from this mean with the sum-to-zero constraint on each of the predictors (∑*μ*_[*predictor*]_ = 0). Table 2 shows the meaning and value of each of the predictors. Generic weakly informative priors were used for all parameters and hyperparameters (*Normal*(0,1)).

**Table 3.**
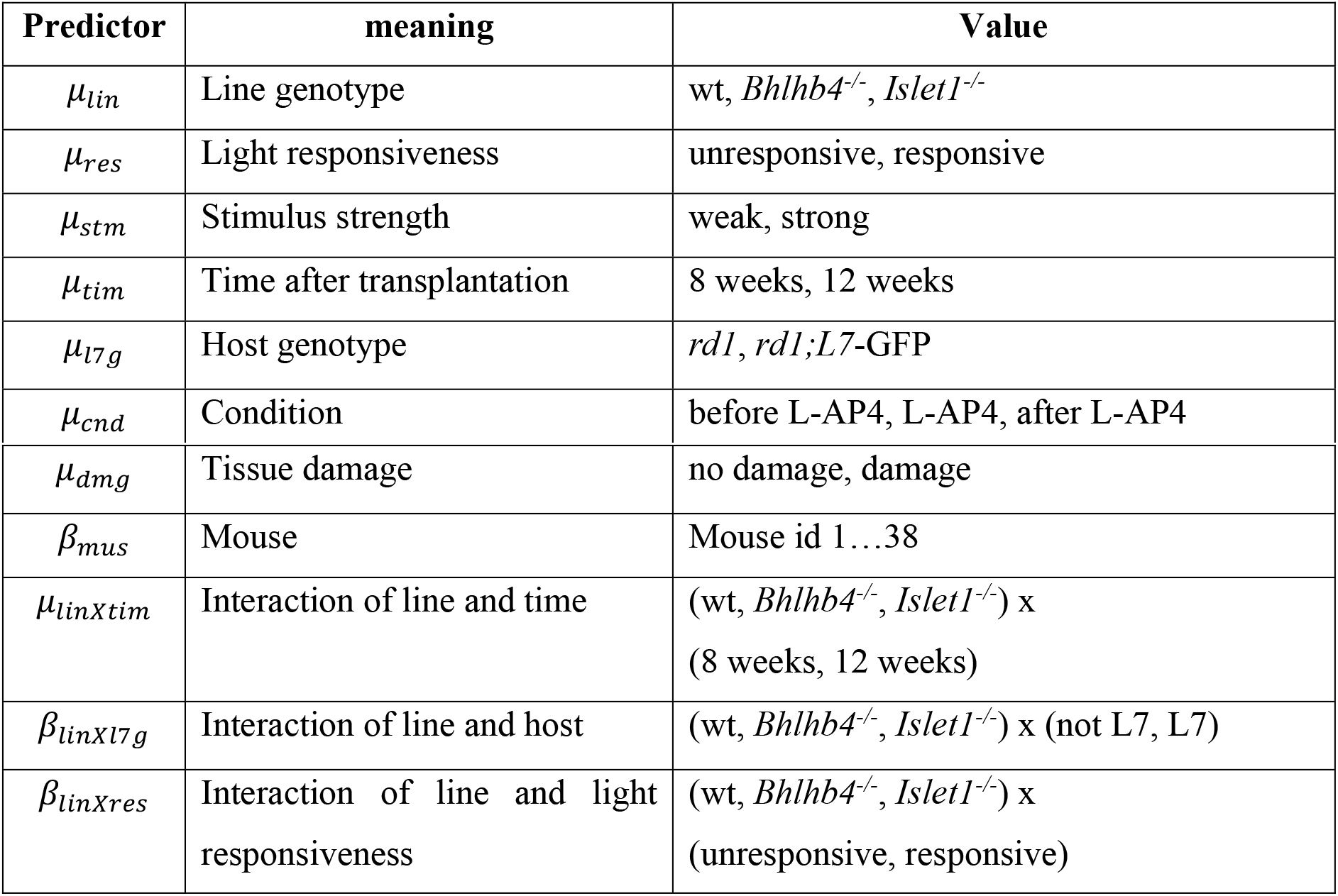
Predictors for analysis of spontaneous activity.

### Behavior: Light Avoidance

The number of successful avoidances *y* out of 30 trials was modeled with the Binomial likelihood.

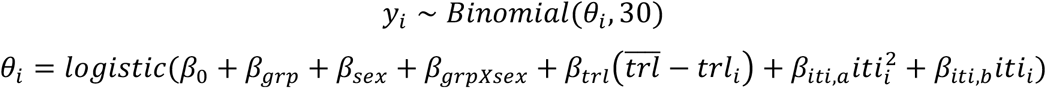

Where *y_i_* represents the number of successful avoidances out of 30 trials and *θ_i_* is the probability of success. *β_grp_* is the effect of different groups (wt, Bhlhb^*-/-*^, *Islet1^-/-^*, and age matched *rd* control without treatment), *β_sex_* is the sex of the animal, and *β_grpXsex_* is the interaction term of sex and group. *β*_0_ represents the overall mean (at mean *trl* and 0 *iti*) and the effect of a particular predictor is modeled as a deviation from this overall mean with a sum-to-zero constraints for each predictor (∑ *β*_[*predictor*]_ = 0). *trl* (trial) is a covariate to account for the accumulated experience, as more experienced mice would be expected to perform better. *iti* is a proxy for the locomotion during the experiment, which greatly influences the success rate as mice randomly moving may avoid the electric shock merely by chance. We assume a quadratic relationship for *iti* rather than linear, however the linear model produced similar results. We believe the quadratic model better describes the data as the random moves of the mouse payoff better at lower *iti* but the benefit of moving faster decreases at high *iti* as the animal tires. Generic weakly informative priors were used for all parameters.

